# BIAFLOWS: A collaborative framework to reproducibly deploy and benchmark bioimage analysis workflows

**DOI:** 10.1101/707489

**Authors:** Ulysse Rubens, Romain Mormont, Lassi Paavolainen, Volker Bäcker, Gino Michiels, Benjamin Pavie, Leandro A. Scholz, Martin Maška, Devrim Ünay, Graeme Ball, Renaud Hoyoux, Rémy Vandaele, Ofra Golani, Anatole Chessel, Stefan G. Stanciu, Natasa Sladoje, Perrine Paul-Gilloteaux, Raphaël Marée, Sébastien Tosi

## Abstract

Automated image analysis has become key to extract quantitative information from scientific microscopy bioimages, but the methods involved are now often so refined that they can no longer be unambiguously described using written protocols. We introduce BIAFLOWS, a software tool with web services and a user interface specifically designed to document, interface, reproducibly deploy, and benchmark image analysis workflows. BIAFLOWS allows image analysis workflows to be compared fairly and shared in a reproducible manner, safeguarding research results and promoting the highest quality standards in bioimage analysis. A curated instance of BIAFLOWS is available online; it is currently populated with 34 workflows that can be triggered to process image datasets illustrating 15 common bioimage analysis problems organized in 9 major classes. As a complete case study, the open benchmarking of 7 nuclei segmentation workflows, including classical and deep learning techniques, was performed on this online instance. All the results presented can be reproduced online.

## Introduction

As life scientists collect microscopy datasets of increasing size and complexity [1], computational methods to extract quantitative information from these images have become inescapable. In turn, modern image analysis methods are becoming so complex (often involving a combination of image processing steps and deep learning techniques) that they require expert configuration and specific execution environments to run. Unfortunately, the software implementations of these methods are commonly shared as poorly reusable and scarcely documented source code, and seldom as user friendly packages for mainstream BioImage Analysis (BIA) platforms [2][3][4]. Even worse, test images are not consistently provided with the software and it can hence be difficult to identify what is the baseline for valid results, or the critical adjustable parameters to optimize the analysis. Altogether, this does not only impair the reusability of the methods and impede reproducing published results [5][6], but it also makes it difficult to adapt these methods to process similar images. To improve this situation, scientific datasets are now increasingly made available through public web-based applications [7][8][9] and open data initiatives [10], but existing platforms do not systematically offer advanced features such as the ability to view and process multidimensional images online, or to let users assess the quality of the analysis against a ground-truth reference (a.k.a. benchmarking). Benchmarking is at the core of biomedical image analysis challenges and it is a practice known to sustain the continuous improvement of BIA (Bio Image Analysis) methods and to promote their wider diffusion [11]. Unfortunately, challenges are rather isolated competitions and they suffer from known limitations [12]: Each event focuses on a single bioimage analysis problem and it relies on ad-hoc data formats and scripts to compute benchmark metrics. Both challenge organizers and participants are therefore duplicating efforts from challenge to challenge, whereas participants’ workflows are rarely available in a sustainable and reusable fashion. Additionally, the vast majority of challenge datasets come from medical imaging, not from biology: for instance, as of January 2020, only 15 out of 198 Grand-challenges datasets [13] were collected from fluorescence microscopy, one of the most common imaging techniques for research in biology. As a consequence, efficient BIA techniques are nowadays available but their reproducible deployment and benchmarking are still stumbling blocks for open science. In practice, end users are hence faced with a plethora of BIA ecosystems and workflows, each with its own specifics, and they have a hard time reproducing other experiments, validating their own analysis, or ensuring that a given technique is the most appropriate for the problem they face. On the other hand, developers cannot systematically validate the performance of their workflow on public datasets, or compare their results to previous work without investing timeconsuming and error prone reimplementation efforts. It is also very challenging for them to make their contribution available to the whole scientific community in a configuration free and safely reproducible manner.

### BIAFLOWS software architecture for reproducible deployment and benchmarking

Within the **N**etwork of **Eu**ropean **B**io**I**mage **A**nalyst**s** (NEUBIAS^1^), an important body of work focuses on channelling the efforts of bioimaging stakeholders (including biologists, bioimage analysts and software developers) to ensure a better characterization of existing bioimage analysis workflows, and to bring these tools to a larger number of scientists. Together, we have envisioned and implemented BIAFLOWS (Fig. 1), *a community-driven, open-source software web platform to reproducibly deploy and benchmark bioimage analysis workflows on annotated multidimensional microscopy data*. Whereas some emerging bioinformatics web platforms [14][15] simply rely on *Dockerized* environments and interactive Python notebooks to process scientific data accessible from public repositories, BIAFLOWS offers a versatile and extensible framework to (i) batch import annotated image datasets and organize them into projects (e.g. BIA problems), (ii) encapsulate versioned BIA workflows regardless of their target BIA platform, (iii) batch process the images, (iv) remotely visualize the images together with workflow results, and (v) automatically assess the performance of the workflows from widely accepted benchmark metrics. All content of a BIAFLOWS instance can be browsed, triggered, and interactively explored from the web interface (Fig. 1). To complement benchmarking results, the output of several workflows can also be visualized simultaneously from synchronized image viewers (Fig. 2).

**Figure 1.**
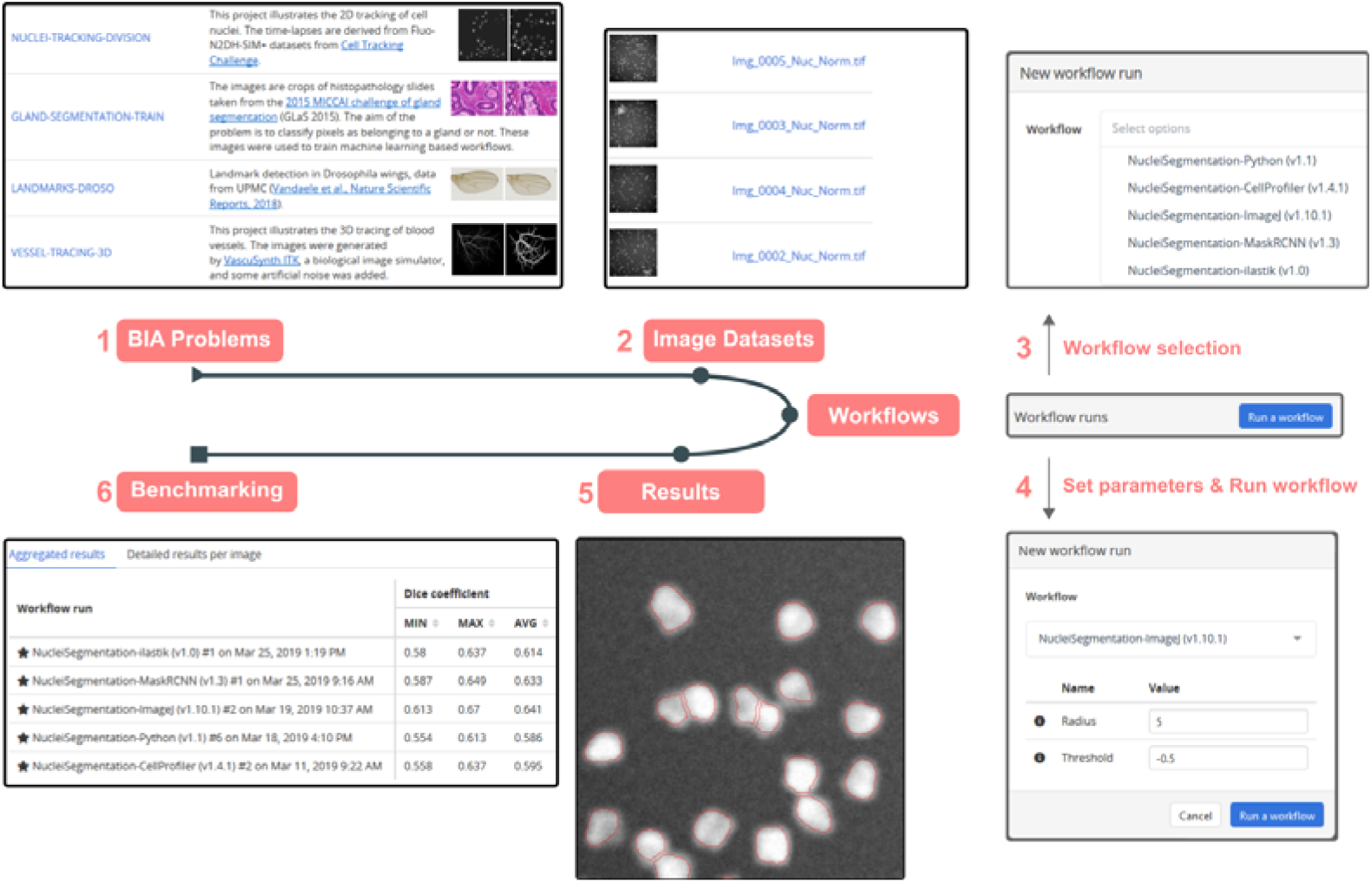
BIAFLOWS web interface. A user selects a problem from the list of BIA problems available from a BIAFLOWS instance (1), browses the images illustrating this problem to compare them to target images (2), and selects a workflow (3) and parameters (4) to process the images. The results of the workflow can then be overlaid on the original images from the online image viewer (5), and benchmark metrics can be browsed, sorted and filtered both as overall statistics (all images), or per image (6).

**Figure 2.**
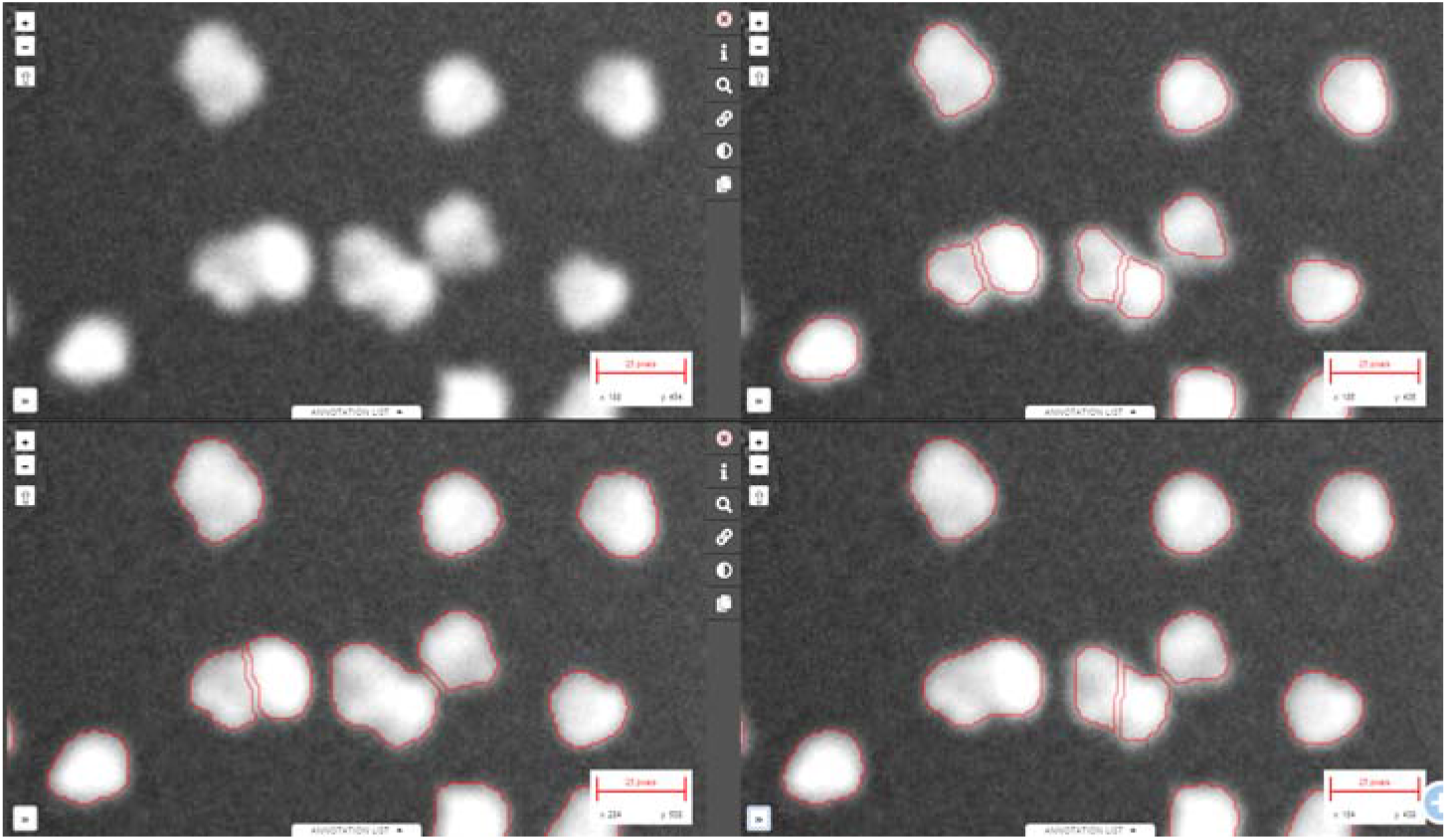
Synchronizing several image viewers displaying the results from several workflow runs. Zoomed in region from one of the images available in NUCLEI-SEGMENTATION problem (accessible from BIAFLOWS online instance). Original image (upper left), same image overlaid with results from: custom ImageJ macro (upper right), custom CellProfiler pipeline (lower left) and custom Python script (lower right).

BIAFLOWS is open-source, thoroughly documented (http://biaflows-doc.neubias.org/) and extends Cytomine [16], a web platform originally developed for the collaborative annotation of high-resolution bright-field bioimages. BIAFLOWS required extensive software developments and content integration to enable the benchmarking of bioimage workflows; accordingly the web user interface has been completely redesigned to streamline this process (Fig. 1). First, a module to upload multidimensional (C, Z, T) microscopy datasets and a full-fledge remote image viewer were implemented. Next, the architecture was refactored to enable the remote execution of bioimage analysis workflows encapsulated with their original software environment in *Docker images* (*workflow images*). To abstract out the operations performed by workflows, we adopted a rich application description schema [17] describing their interface (input, output, parameters) (Supplementary Methods, Section 4). The system was also engineered to monitor trusted user spaces hosting a collection of *workflow images* and to automatically pull new or updated workflows (Fig. 3, DockerHub). In turn, *workflow images* are built and versioned in the cloud whenever a new release is triggered from their associated source code repositories (Fig. 3, GitHub). To ensure reproducibility, we enforced that all versions of the *workflow images* are permanently accessible from the system. Importantly, the workflows can be seamlessly run on any computational resource, including high-performance computing and multiple server architectures. This is achieved by converting the *workflow images* to a compatible format (Singularity, [18]), and dispatching them to the target computational resources over the network (using e.g. SLURM, [19], Fig. 3, additional computing servers). To enable interoperability between all components, standard object annotation formats were specified for 9 important classes of BIA problems (Supplementary Methods, Section 5), and we developed a software library required to compute relevant benchmark metrics associated to these problem classes (partly adapted from biomedical challenges [13] and scientific publications [21]). With this new design, benchmark metrics are automatically computed after each workflow run. BIAFLOWS can also be deployed on a local server to manage private images and workflows, and to process image locally (Fig. 3 BIAFLOWS Local, Supplementary Methods, Section 3). To simplify the coexistence of these different deployment scenarios, migration tools were developed (Supplementary Methods, Section 6) to transfer content between existing BIAFLOWS instances (including the online instance described hereafter). Importantly, provided a user has the corresponding rights, all content from any instance can be accessed programmatically through a RESTful interface, which ensures a complete data accessibility and interoperability. Finally, for full flexibility, workflows can also be downloaded manually from DockerHub to process local images independently of any BIAFLOWS instance (Fig. 3, Standalone Local, Supplementary Methods, Section 6).

**Figure 3.**
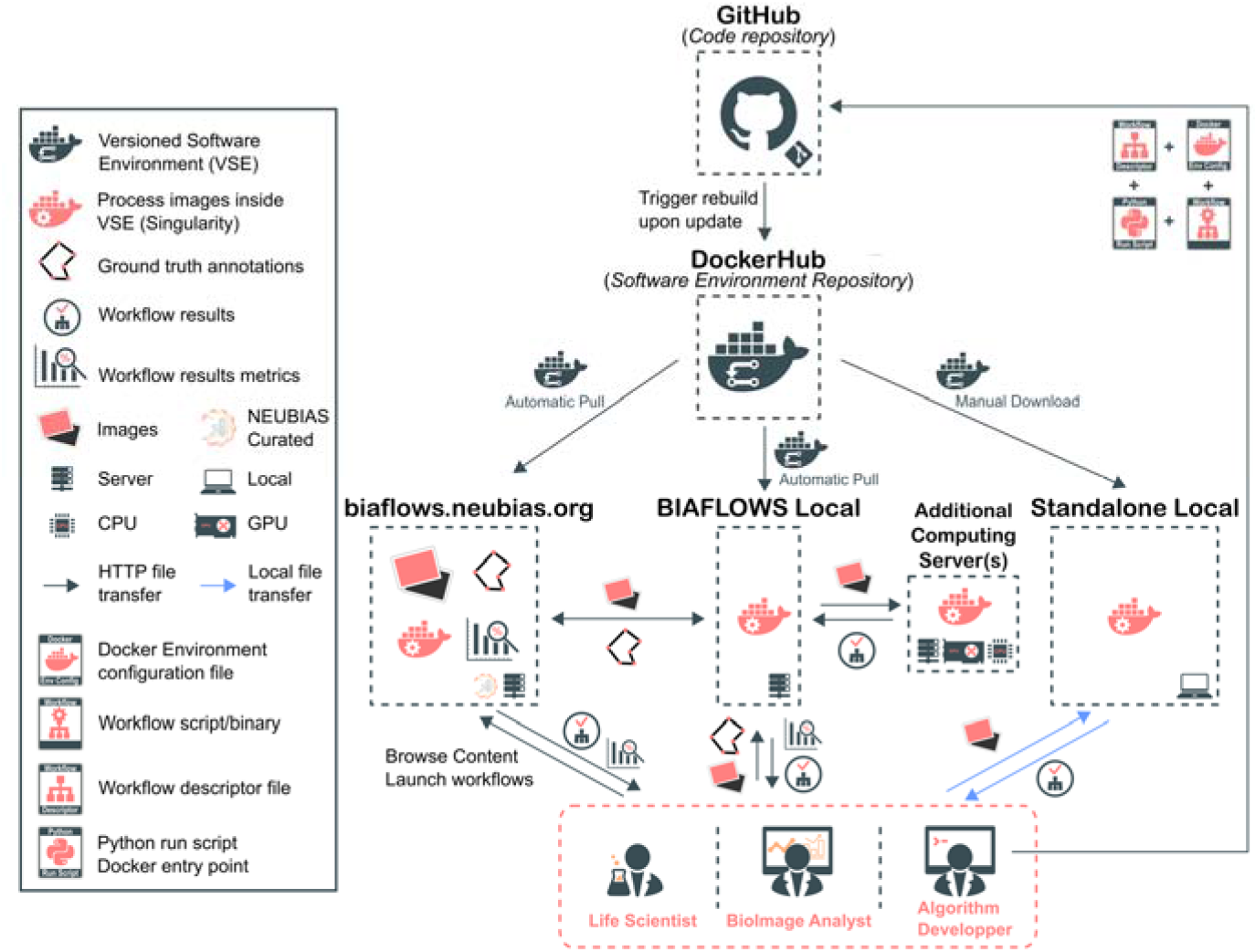
BIAFLOWS architecture and possible deployment scenarios. Workflows are hosted in a trusted source code repository (GitHub). Workflow images (Docker images) encapsulate workflows together with their execution environments to ensure reproducibility. The workflow images are automatically built by DockerHub (cloud service) whenever a new workflow is released or an existing workflow is updated from its trusted GitHub repository. BIAFLOWS instances monitor DockerHub and pull new or updated workflow images. Workflow images can also be downloaded manually to process local images in a standalone fashion.

### BIAFLOWS online, curated, instance for public benchmarking

An online instance of BIAFLOWS is managed by NEUBIAS and available at: https://biaflows.neubias.org/ (Fig. 3). This instance is ready to host community contributions and it is already populated with a substantial collection of *publicly shared* annotated image datasets illustrating important BIA problems, and associated versioned workflows targeting common BIA platforms (Supplementary Methods, Section 1, Table 1). More precisely, we integrated heterogeneous workflows addressing 9 important classes of BIA problems, illustrated by 15 image datasets imported from challenges (DIADEM [22], Cell Tracking Challenge [23], Particle Tracking Challenge [24], Kaggle Data Science Bowl 2018 [25]), created from synthetic data generators [20] (CytoPacq [26], TREES toolbox [27], Vascusynth [28], SIMCEP [29]), or contributed by NEUBIAS members [38]. The following problem classes are currently represented: object detection/counting, object segmentation, and pixel classification (Fig. 4); particle tracking, object tracking, filament network tracing, filament tree tracing and landmark detection (Fig. 5). To demonstrate the versatility of the platform, 34 heterogeneous BIA workflows were integrated, and each target a specific BIA platform, language, or library: ImageJ/FIJI macros and scripts [30], Icy protocols [31], CellProfiler pipelines [32], Vaa3D plugins [33], ilastik pipelines [34], Octave scripts [40], Jupyter notebooks [15], and Python scripts leveraging Scikit-learn [35] for supervised learning algorithms, and Keras/PyTorch [36][37] for deep learning. This list, although already extensive, is not limited, as BIAFLOWS core architecture enables to seamlessly integrate other software environments as long as the workflows check minimalistic requirements (Supplementary Methods, Section 4). So far, the workflows integrated to the platform were mostly contributed by scientists from NEUBIAS Workgroup 5, or imported from existing challenges by reproducibly encapsulating original author’s code into their specified execution environments. In order to demonstrate the potential of BIAFLOWS to perform open benchmarking, a complete case study has been performed with (and is available from) BIAFLOWS to compare workflows segmenting nuclei in heterogeneous 2D microscopy images (Supplementary Methods, Section 2). The content from BIAFLOWS online instance (https://biaflows.neubias.org) can be viewed in read only mode from the guest account (user: *guest* | password: *guest*). Please contact us if you want to get workflow execution rights. An extensive user guide and video tutorial are available online from the same URL. To enhance the visibility of the workflows hosted in the system, all workflows are referenced from NEUBIAS Bioimage Informatics Search Index (http://biii.eu/). The online instance is fully extensible and, with minimal efforts, interested developers can package their own workflows (Supplementary Methods, Section 4) and make them available for benchmarking from BIAFLOWS online instance (see Box 2. How to contribute). Similarly, following our guidelines (Supplementary Methods, Section 3), scientists can make their images and ground-truth annotations available online through BIAFLOWS online instance or a BIAFLOWS instance they would manage locally (see Box 2. How to contribute). All content from the online instance can be seamlessly migrated to a local BIAFLOWS instance (Supplementary Methods, Section 6) for further development or to process local images.

**Figure 4.**
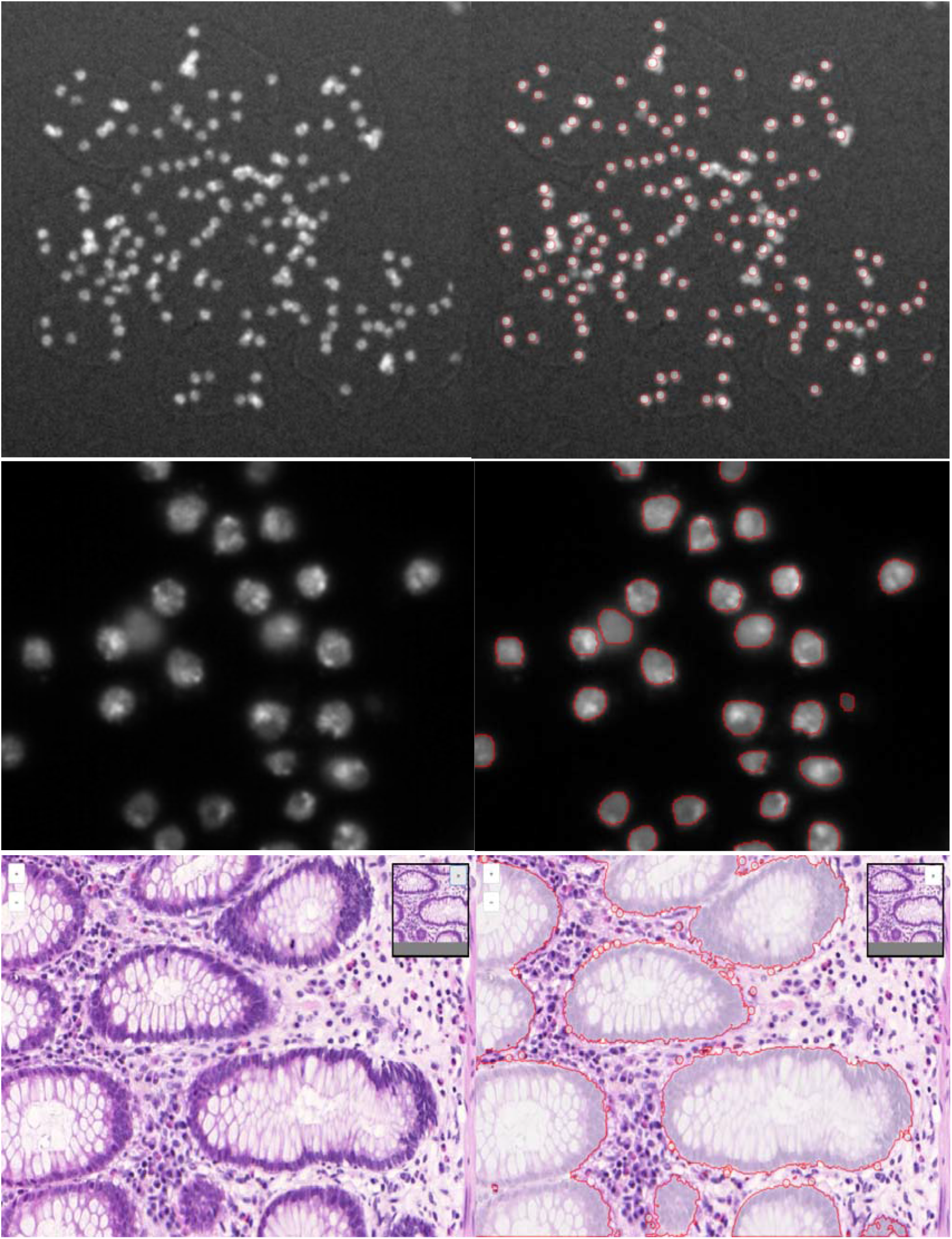
Some sample images from BIAFLOWS online instance illustrating different BIA problem classes, and the results from an associated workflow. Original image (left) and workflow results (right), from top to bottom: 1. Spot / object detection in synthetic image displaying spots (SIMCEP [38]). 2. Nuclei segmentation in images from Data Science Bowl 2018 [25]. 3. Pixel classification in images from 2015 MICCAI gland segmentation challenge [38].

**Figure 5.**
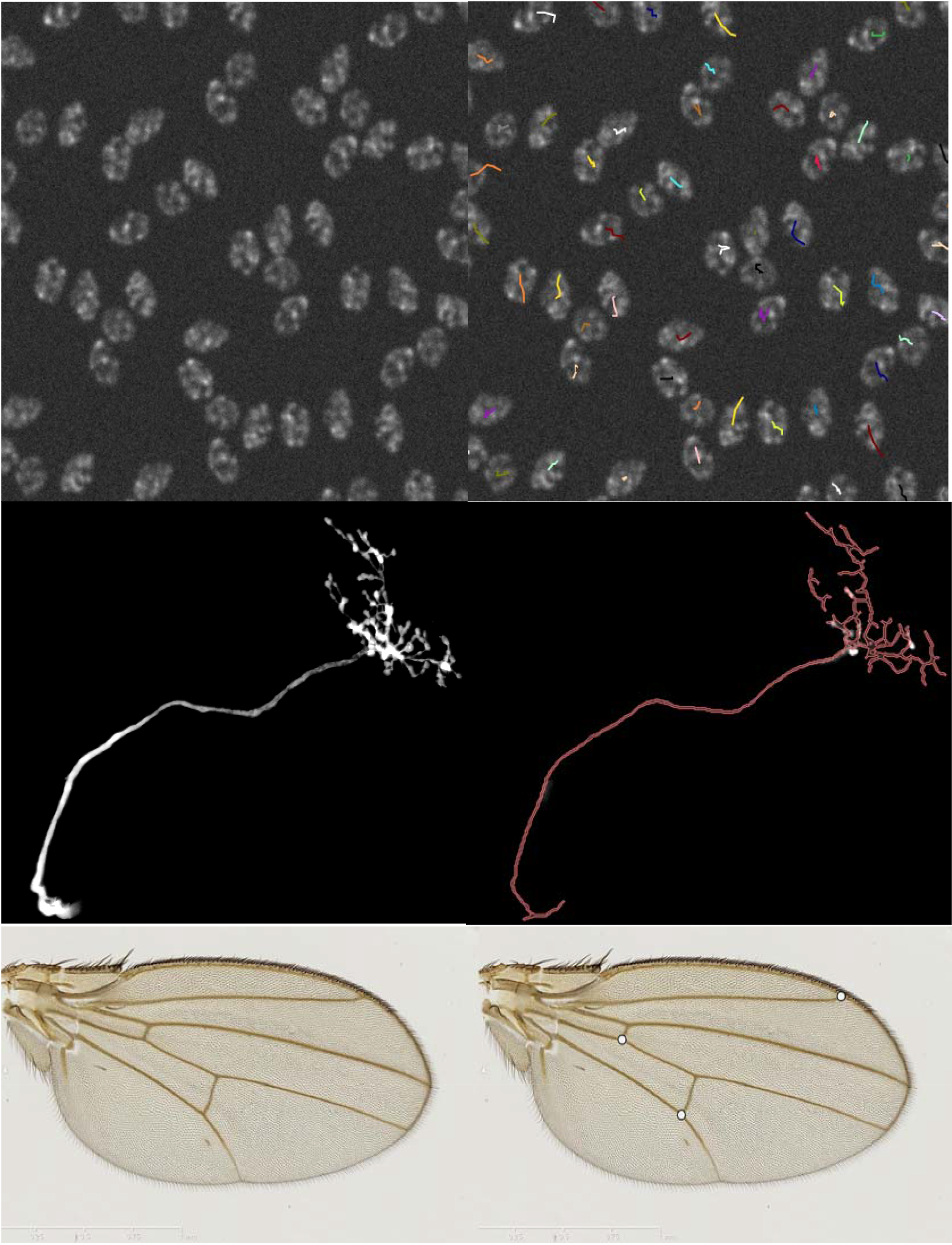
Some sample images from BIAFLOWS online instance illustrating different BIA problem classes, and the results from an associated workflow. Original image (left) and workflow results (right), from top to bottom: 1. Particle tracking in synthetic timelapse displaying non-dividing nuclei (CytoPACQ [25]), single frame + dragon tail tracks. 2. Neuron tree tracing in 3D image stacks from DIADEM challenge [21], average intensity projection (left), traced skeleton dilated Z-projection (red). 3. Landmark detection in Drosophila wing images [37].

#### Box 1. How to get started with BIAFLOWS

- Watch BIAFLOWS video tutorial (https://biaflows.neubias.org)
- Visit BIAFLOWS documentation portal (http://biaflows-doc.neubias.org), including a detailed user guide and instructions for developers and data contributors.
- Access BIAFLOWS online instance (https://biaflows.neubias.org) in read-only mode from the guest account (username: *guest* password: *guest*). The platform is maintained by NEUBIAS (http://neubias.org) and backed by bioimage analysts and software developers across the world.
- Install your own BIAFLOWS instance on a desktop computer or a server to manage your images locally or process them with existing BIAFLOWS workflows. Follow *Installing and populating BIAFLOWS locally* from the documentation portal.
- Use an existing workflow to process your own images locally. Follow *Executing a BIAFLOWS workflow BIAFLOWS server* from the documentation portal.
- Share your thoughts or get help on the image.sc forum (https://forum.image.sc), or write directly to our developer team.

#### Box 2. How to contribute to BIAFLOWS

It is possible to contribute to BIAFLOWS in many ways:

- Scientists can contribute annotated images representative of a specific experiment to BIAFLOWS online instance (send a request to), or setup their own BIAFLOWS instance to share these images. See *Problem classes, ground truth annotations and reported metrics* from the documentation portal for guidelines on how to format the ground truth annotations of your images.
- Workflow developers can encapsulate their code following our developer guidelines, test them on a local BIAFLOWS instance on public images migrated to their local server (using our migration tool), or publish their source code on GitHub to showcase their workflows in BIAFLOWS online instance. Follow *Creating a BIA workflow and adding it to a BIAFLOWS instance* from the documentation portal and submit contributions to.
- Feature requests or bug reports can be posted to BIAFLOWS GitHub (https://github.com/neubias-wg5).
- Share your experience with us and others, help other researchers on the image.sc forum (https://forum.image.sc).
- Contribute to the documentation by submitting a pull request to our documentation repository (https://github.com/Neubias-WG5/neubias-wg5.github.io).
- Cite and link BIAFLOWS content to share data and results accompanying your scientific publications. Follow *Access BIAFLOWS from a Jupyter notebook* from the documentation portal, or directly link the webpages of your BIAFLOWS instance.

To further increase the current content of BIAFLOWS online instance, calls for contribution will be launched to the scientific community to gather more microscopy annotated images and encourage developers to package their own workflows (see Box 2. How to contribute). The support of new problem classes is also planned, e.g. to benchmark the detection of blinking events in the context of super-resolution localization microscopy or the detection of landmark points for image registration. There is actually no limitation in using BIAFLOWS in other fields where image analysis is a critical step to extract scientific results from images, for instance material science and biomedical imaging.

### Case study: Benchmarking nuclei segmentation by classical image processing, machine learning, and deep learning methods

We integrated 7 nuclei segmentation workflows to BIAFLOWS online instance and used the platform for their open benchmarking (Supplementary Methods, Section 2). All content of this case study (images, ground-truth annotations, workflow codes and software environment, workflow results) are readily accessible from BIAFLOWS online instance (see Supplementary Methods, Section 2). The workflows were benchmarked on two different image datasets: a synthetic dataset made of 10 images generated for the purpose of this case study, and a subset of 65 images coming from an existing nuclei segmentation challenge (Science Bowl 2018). In this study, we could identify trends among the different techniques implemented by the workflows: (i) optimal image pre-processing can bring an edge to object segmentation accuracy, even when classical techniques compete with deep learning techniques (ii) workflows relying on deep learning achieve better than classical workflows for heterogeneous datasets, and (iii) object de-clumping is a critical postprocessing step for object segmentation accuracy. It was also evidenced that, for meaningful comparison, a set of benchmarking metrics is to be favoured to a single metric (e.g. DICE coefficient), and that visual inspection of the workflow results is often required to understand the actual shortcomings highlighted by poorer metric scores. All these features are enabled by BIAFLOWS, and the same methodology applied in this case study can be easily translated to other benchmarking studies.

## Conclusion and future developments

BIAFLOWS addresses a number of critical requirements to foster open image analysis for Life Sciences: (i) sharing and visualizing annotated images illustrating commonly faced BIA problems, (ii) sharing versioned, reproducible, BIA workflows, (iii) exposing important workflow parameters and their sensible default values, (iv) computing relevant performance metrics to compare the workflows, and (v) providing a standard way to store, visualize, and share the results of the workflows. As such, BIAFLOWS is a critical asset for biologists and bioimage analysts to leverage state of the art bioimage techniques and efficiently reuse them to process their own data. It is also a tool of choice for algorithm developers and challenge organizers to benchmark bioimage analysis workflows and identify trends among different analysis strategies. For challenges, some BIAFLOWS projects would be made public to readily provide training datasets to competitors, while others would remain private and would be the ones used to benchmark the competing workflows. Finally, BIAFLOWS can help authors willing to share online supporting data, methods and results associated to their published scientific results. More generally, the platform could be adopted as a central and federated platform to make scientific images available to the community and publish them together with the workflows that were used to extract annotations from these images. This is a milestone of open science not reached yet by any image sharing platform and it should help accelerating scientific progress and surpassing individualist practices where image datasets, algorithms, quantification results, and associated knowledge are often stored locally or with restricted access.

## Acknowledgments

This project is funded by COST CA15124 (NEUBIAS). BIAFLOWS is developed by NEUBIAS (http://neubias.org) Workgroup 5 and it would not have been possible without the great support from NEUBIAS vibrant community of bioimage analysts, and the dedication of Julien Colombelli and Kota Miura in organizing this network. Local organizers of the Taggathons who have fostered the development of BIAFLOWS are also greatly acknowledged: Chong Zhang, Gabriel G. Martins, Julia Fernandez-Rodriguez, Peter Horvath, Bertrand Vernay, Aymeric Fouquier d’Hérouël, Andreas Girod, Paula Sampaio. We also thank the various developers who answered our questions regarding the integration of their code, among others the Image Analysis Hub from Pasteur Institute. **RV** was supported by ADRIC Pôle Mecatech Wallonia grant, and **UR** by ADRIC Pôle Mecatech and DeepSport Wallonia grants. **RaM** was supported by IDEES grant with the help of the Wallonia and the European Regional Development Fund (ERDF). **LP** was supported by Academy of Finland (grant 310552). **MM** was supported by the Czech Ministry of Education, Youth and Sports (project LTC17016). **SGS** acknowledges the financial support of UEFISCDI grant PN-III-P1-1.1-TE-2016-2147 (CORIMAG). **VB** and **PPG** acknowledge the France-BioImaging infrastructure supported by the French National Research Agency (ANR-10-INBS-04).

## Author contribution

**ST** and **RaM** conceptualized BIAFLOWS, supervised its implementation, contributed to all technical tasks and wrote the manuscript. **UR** worked on the core implementation of BIAFLOWS with contributions from **GM** and **RH**. **RoM** implemented several modules to interface bioimage analysis workflows and the content of the system. **ST, MM** and **DU** implemented the module to compute benchmark metrics. **ST**, **VB**, **RoM**, **LP**, **BP**, **MM, RV**, and **LAS** integrated their own workflows or adapted existing workflows, and tested the system. **ST**, **DU**, **OG,** and **GB** organized and collected content (image datasets, simulation tools). **SGS**, **NS** and **PP-G** provided extensive feedback on BIAFLOWS and contributed to the manuscript. All authors took part in reviewing and improving the manuscript.

## Data availability

All images and annotations can be downloaded from the online instance of BIAFLOWS at https://biaflows.neubias.org/.

## Code availability

BIAFLOWS is an open source project and its code source can be downloaded at https://github.com/Neubias-WG5.

## Documentation

http://biaflows-doc.neubias.org/

## Footnotes

1 NEUBIAS: Network of European BioImage Analysts, COST (http://www.cost.eu) Action CA15124 (NEUBIAS)

