## Supplementary material for "BIAFLOWS: A collaborative framework to reproducibly deploy and benchmark bioimage analysis workflows"

**Ulysse Rubens**° (Université de Liège), **Romain Mormont**° (Université de Liège), Lassi Paavolainen (FIMM, HiLIFE, University of Helsinki), Volker Bäcker (MRI, BioCampus Montpellier), Gino Michiels (HEPL / Université de Liège), Benjamin Pavie (VIB BioImageing Core), Leandro A. Scholz (Universidade Federal do Paraná), Martin Maška (Masaryk University), Devrim Ünay (Ízmir University of Economics), Graeme Ball (Dundee Imaging Facility, School of Life Sciences, University of Dundee), Renaud Hoyoux (Cytomine SCRL FS), Rémy Vandaele (Université de Liège), Ofra Golani (Life Sciences Core Facilities, Weizmann Institute of Science), Anatole Chessel (École Polytechnique), Stefan G. Stanciu (Politehnica Bucarest), Natasa Sladoje (Uppsala University), Perrine Paul-Gilloteaux (Structure Fédérative de Recherche François Bonamy, Université de Nantes, CNRS, INSERM), **Raphaël Marée*** (Université de Liège), **Sébastien Tosi*** (Institute for Research in Biomedicine, IRB Barcelona, Barcelona Institute of Science and Technology, BIST)

° These authors contributed equally to this work

* These authors supervised this work (+ corresponding authors)

### Supplementary Section 1. Content of BIAFLOWS online instance

Table 1 reports the BIA problems and workflows currently available from BIAFLOWS online instance, and that can be imported from Github to any other BIAFLOWS instance. It consists of 34 workflows spanning 15 BIA problems (from 9 different classes).

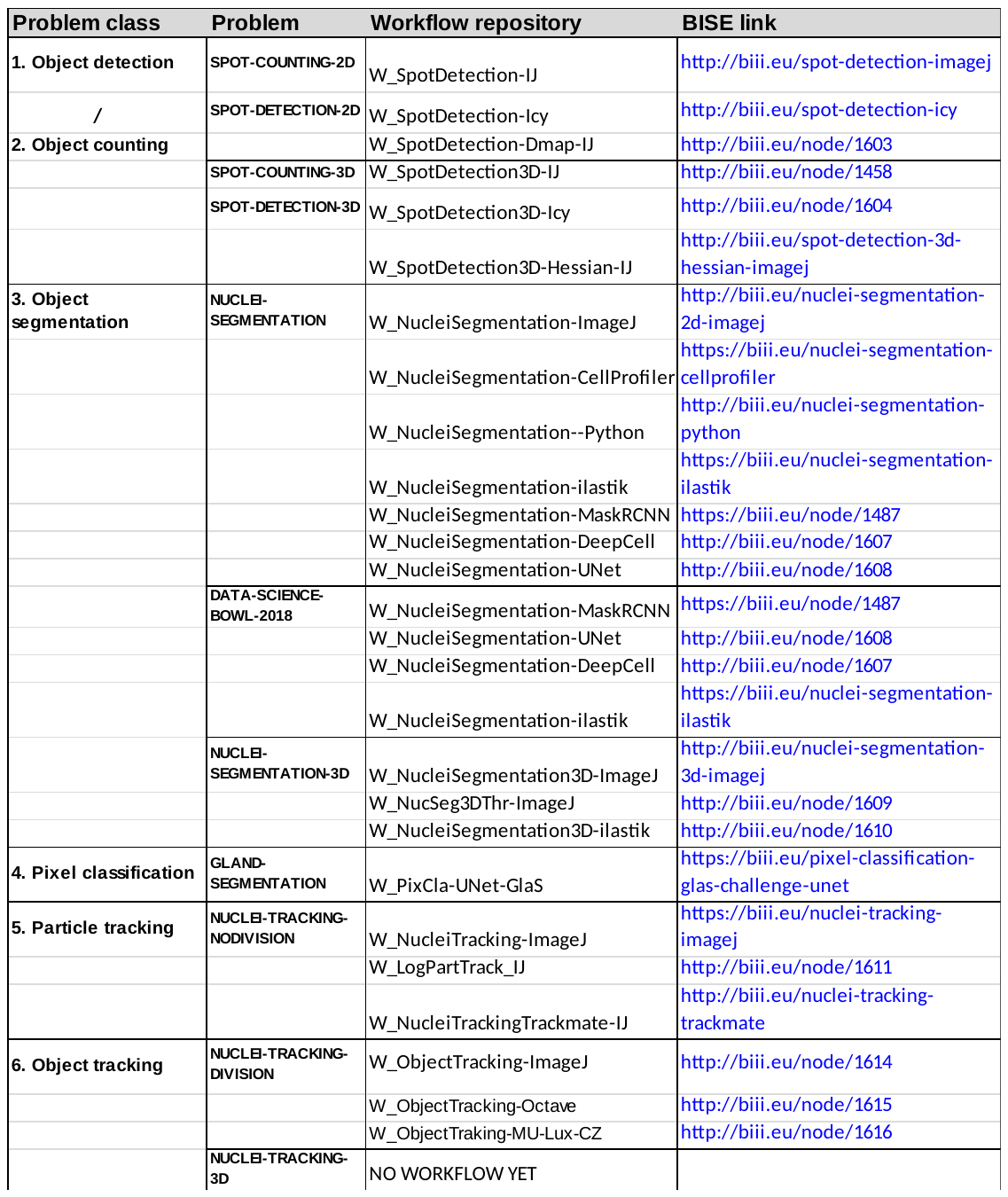

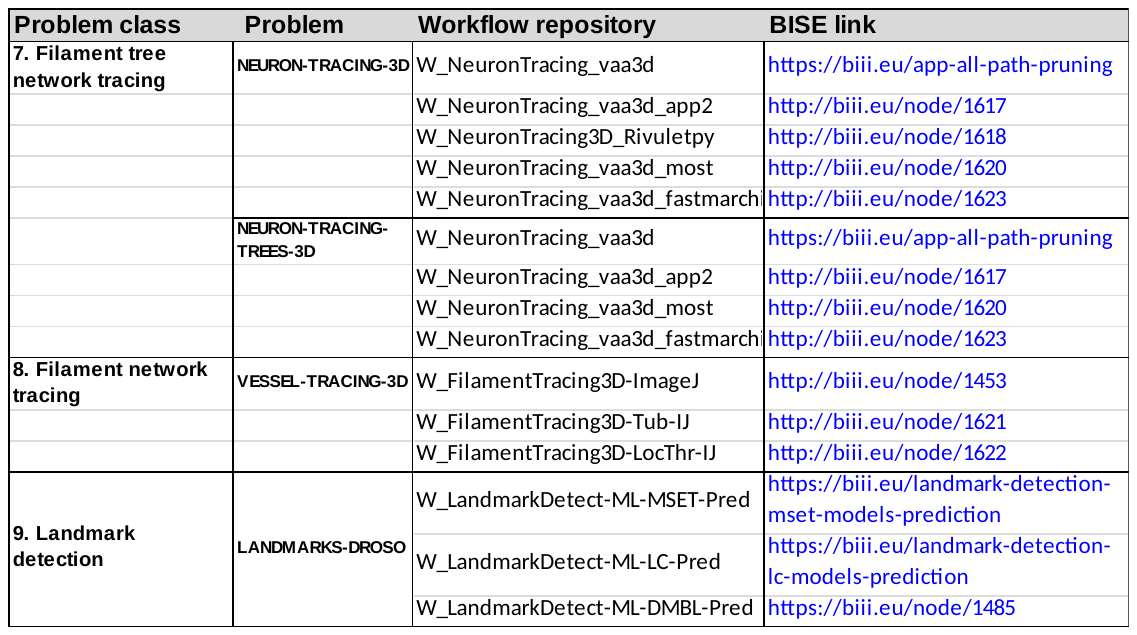

**Supplementary Table 1. BIA problems and workflows currently available from BIAFLOWS online instance.** **Problem**: BIA problem as listed from BIAFLOWS > Problems webpage. **Problem class**: the type of image analysis. **Workflows repository**: the name of the workflow source code repository as linked from BIAFLOWS > Workflows webpage (all repositories are stored on <https://github.com/Neubias-WG5> GitHub). **BISE link**: the workflow reference webpage on BISE, NEUBIAS Bioimage Informatics Search Engine (biii.eu).

### Supplementary Section 2. Benchmarking case study: nuclei segmentation (2D images)

In this Section, we provide details for the benchmarking case study and web links to access all data.

**Datasets**

1. [**BIAFLOWS NUCLEI-SEGMENTATION**](https://biaflows.neubias.org/#/project/5955/images) **(SIMCEP)**

10 synthetic grayscale images simulating widefield fluorescence microscopy images (SIMCEP simulator). The images suffer from strong non-uniform illumination, half of them are heavily saturated, and nuclei are largely clustered in some regions.

2. [**BIAFLOWS DATA-SCIENCE-BOWL-2018**](https://biaflows.neubias.org/#/project/12182234/images) **(DSB)**

65 RGB images taken from Data Science Bowl 2018 challenge and representing nuclei stained from heterogeneous samples imaged by different microscopy modalities.

**Workflows**

| **Name** | **Pre-processing** | **Classification** | **Mask post-processing** |
| --- | --- | --- | --- |
| [ImageJ](https://github.com/Neubias-WG5/W_NucleiSegmentation-ImageJ) | Laplacian of Gaussian | Global threshold  (user defined) | Binary watershed from distance map local maxima |
| [Python](https://github.com/Neubias-WG5/W_NucleiSegmentation-Python) | Gaussian blur | Adaptive threshold  (local mean) | Binary watershed from distance map regional maxima |
| [CellProfiler](https://github.com/Neubias-WG5/W_NucleiSegmentation-CellProfiler) | Gaussian blur and illumination  correction | Global threshold  (Otsu’s method) | De-clumping from Laplacian of Gaussian local minima |
| [Ilastik](https://github.com/Neubias-WG5/W_NucleiSegmentation-ilastik)1* | None | Foreground / background feature based pixel classifier^[1]^. Trained on SIMCEP images from scarce hand annotations. | Remove small objects |
| [Ilastik](https://github.com/Neubias-WG5/W_NucleiSegmentation-ilastik)2* | None | Foreground / background feature based pixel classifier^1^. Trained on DSB images from scarce hand annotations. | Remove small objects |
| [Mask-RCNN](https://github.com/Neubias-WG5/W_NucleiSegmentation-MaskRCNN)* [1] | None | Foreground / background classifier trained^[2]^ on 670 images from DSB2018 training set 1. | None, but classifier accounts for object geometry |
| [U-NET](https://github.com/Neubias-WG5/W_NucleiSegmentation-UNet)* [2] | None | Background / edge / interior classifier trained^[3]^ as in [4] on 670 images from DSB2018 training set 1. | Fill holes, remove small objects |
| [DeepCell1.0](https://github.com/Neubias-WG5/W_NucleiSegmentation-DeepCell)* [3] | None | Background / edge / interior classifier (trained model from [3]). Training was performed from BBBC039 [6], a subset of 200 images from DSB2018. | Fill holes, remove small objects |

**Metrics**

**Dice coefficient (DICE, 0-1)**: normalized overlap between ground truth and prediction binary masks (0: no overlap, 1: perfect overlap).

**Average Hausdorff Distance (AHD, >=0)**: average distance between object pixels in ground truth (/prediction) masks and closest object pixels in prediction (/ground truth) masks.

**Fraction overlap (FO, 0-1)**: 0.5 fraction overlap can be interpreted as "on average the overlap of a predicted object with the ground truth object with largest overlap is half the area of the larger of these two objects". This would for instance happen if objects are either systematically split into two identical objects or merged by pair.

**Mean Average Precision (mAP, 0-1) [4]:** IoU (Intersection of Union) between predicted and ground-truth objects are computed. IoU are then compared to 10 thresholds (0.5, 0.55 ... 0.95) and, if greater, the object is set as true positive (TP) for that threshold. Precision is computed as P = TP / (TP + FP + FN) for each threshold, where FP = number of predicted objects - TP and FN = number of ground-truth objects - TP. Precision is finally averaged out for all objects and images.

**Results**

**NUCLEI-SEGMENTATION dataset**

All results, including workflow parameter values, metrics, and detected nuclei within images are available online: [**https://biaflows.neubias.org/#/project/5955/analysis**](https://biaflows.neubias.org/#/project/5955/analysis) and discussed below.

For this dataset, U-NET and DeepCell are the most accurate workflows to classify nuclei pixels (Supplementary Table 2, DICE, AHD). From visual inspection, Mask-RCNN poorer AHD likely stems from missed nuclei bits (Figure 1.c). Python and CellProfiler workflows achieve the worse DICE; probably since they both implement Gaussian blur pre-processing (Figure 1.a, blurring leads to spatial overestimation of the nuclei), while Python workflow achieves the worse AHD: ilastik1 workflow misses nuclei bits (Figure 1.b); Python workflow overestimates nuclei (Figure 1.a). Finally, CellProfiler workflow achieves a better balance and ImageJ achieves an excellent score for these two metrics (close to best results).

|  | **DICE** | **AHD** | **FO** | **MAP** |
| --- | --- | --- | --- | --- |
| **ImageJ** | 0.912 | 0.119 | **0.821** | **0.593** |
| **Mask-RCNN** | 0.896 | 0.242 | 0.769 | 0.555 |
| **U-NET** | **0.927** | 0.095 | 0.748 | 0.522 |
| **DeepCell** | 0.923 | **0.087** | 0.719 | 0.481 |
| **CellProfiler** | 0.866 | 0.183 | 0.709 | 0.453 |
| **Python** | 0.852 | 0.467 | 0.677 | 0.402 |
| **Ilastik1** | 0.89 | 0.23 | 0.696 | 0.166 |

**Supplementary Table 2.** **Benchmarking results for the 7 2D nuclei segmentation workflows**. For each metric, best results in bold, green highlights results significantly better than average, and orange / red highlight results worse / significantly worse than average.

ImageJ workflow is the best achieving workflow for both FO and MAP (Figure 1.f). This shows that Laplacian of Gaussian filter is an efficient pre-processing step to segment blobs of similar size over non uniform background. Interestingly, Mask-RCNN achieves second despite that it was trained on a different dataset.  U-NET and DeepCell both achieve worse FO and MAP, probably since they tend to merge touching objects (Figure 1.d). Finally, ilastik1 is the worse achieving workflow for both metrics, which clearly correlates with the visual results: many touching nuclei are merged and nuclei are fuzzy with frequent holes (Figure 1.b). This is partly explained by the fact that ilastik workflow does not implement any object splitting post-processing step. Also, it was difficult to accurately train ilastik classifier for both non saturated and saturated images. To this respect, deep learning algorithms are way more flexible. Python workflow exhibits the worse FO, probably reflecting nuclei overestimation. CellProfiler achieves slightly better for both metrics, its most frequent issue is to erroneously merge single nuclei (Figure 1.e).

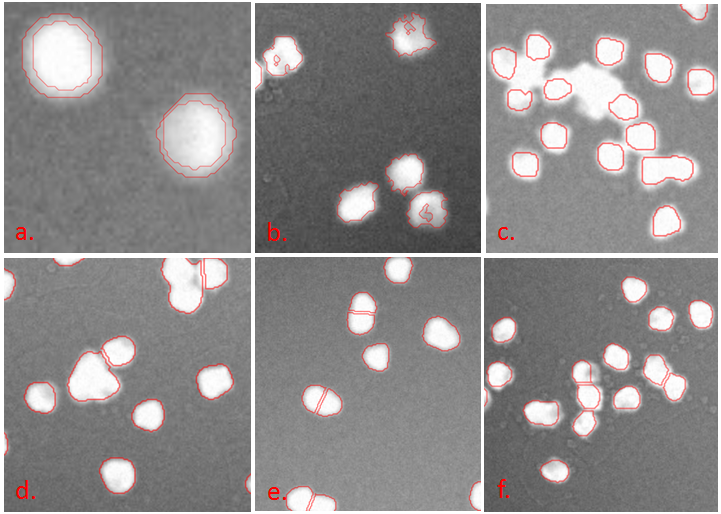

**Figure 1.** **Some snapshots illustrating typical segmentation errors.** **a**. Python workflow (outer rings) + reference nuclei, **b**. ilastik workflow, **c**. Mask-RCNN workflow, **d**. CellProfiler workflow, **e**. Python workflow, **f**. ImageJ workflow.

**Data Science Bowl 2018 dataset**

All results, including workflow parameter values, metrics, and detected nuclei within images are available online: [**https://biaflows.neubias.org/#/project/12182234/analysis**](https://biaflows.neubias.org/#/project/12182234/analysis) and discussed below.

Overall, all metrics are significantly worse than for the previous dataset (Supplementary Table 2), showing that this dataset is more challenging. Classic workflows were not assessed since they are not adapted to non-heterogeneous RGB images. All machine learning workflows perform reasonably close, except ilastik2 workflow which displays significantly worse FO and MAP. From visual inspection, Mask-RCNN does the best job at splitting touching objects but it misses some objects. U-NET is more sensitive (better DICE) but it merges many touching objects. mAP results are significantly lower than the best workflows competing in DSB 2018 (~0.4) since no post-processing was performed on the masks (e.g. object declumping).

|  | **DICE** | **AHD** | **FO** | **mAP** |
| --- | --- | --- | --- | --- |
| **U-NET** | **0.783** | 6.39 | **0.559** | **0.284** |
| **Mask-RCNN** | 0.739 | **2.78** | 0.553 | 0.27 |
| **Ilastik2** | 0.748 | 3.201 | 0.472 | 0.105 |

**Supplementary Table 3.** **Benchmarking results for the 4 machine learning workflows**. For each metric, best results in bold.

**Discussion**

This benchmarking test case stresses the need to use a set of benchmarking metrics as opposed to a single metrics (e.g. the ubiquitous DICE coefficient), and it confirms the versatility of deep learning algorithms to deal with heterogeneous datasets, as observed by [4]. Overall, FO and MAP correlate very well with visual inspection, MAP being more sensitive than the simpler FO metric. DICE and AHD, while informative, are not sufficient to assess object segmentation accuracy since they do not account for erroneous object splitting / merging events. Importantly, even though the selected set of metrics fairly captures the actual performance of the workflows, visually inspecting the results was important to conclude on the actual shortcomings of the workflows. This feature and the ability to browse benchmarking results both as agglomerated statistics and per image are both natively supported by BIAFLOWS. All results are publically available and can be reproduced online, which contrasts with common benchmarking publication practices, e.g. [4], where the code provided requires expert setup to run locally. Figures 2. To 7. illustrate BIAFLOWS capability to explore and share data and results online.

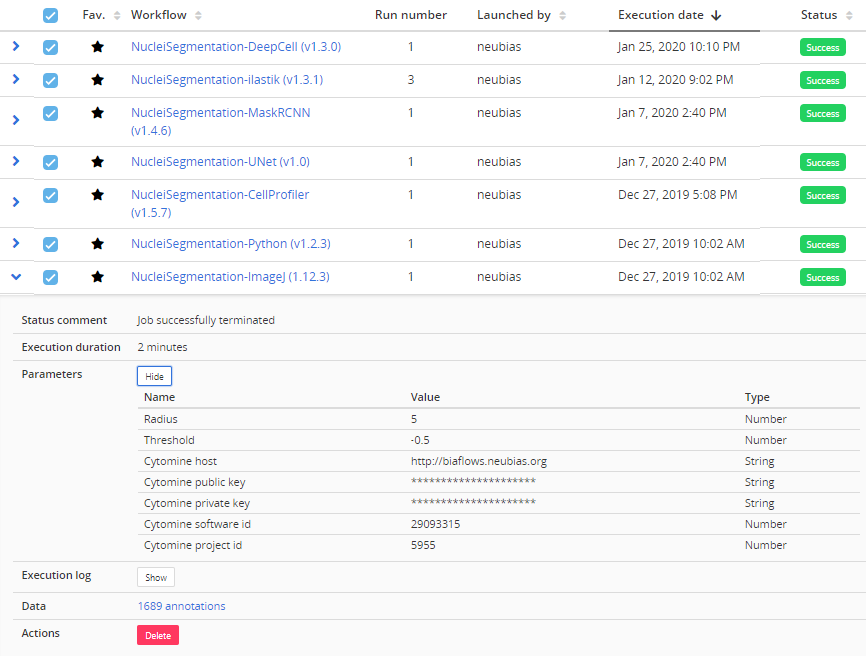

**Figure 2. BIAFLOWS benchmarked workflow runs for SIMCEP dataset.** For every run, the parameters used and the execution log can be retrieved from drop down tabs.

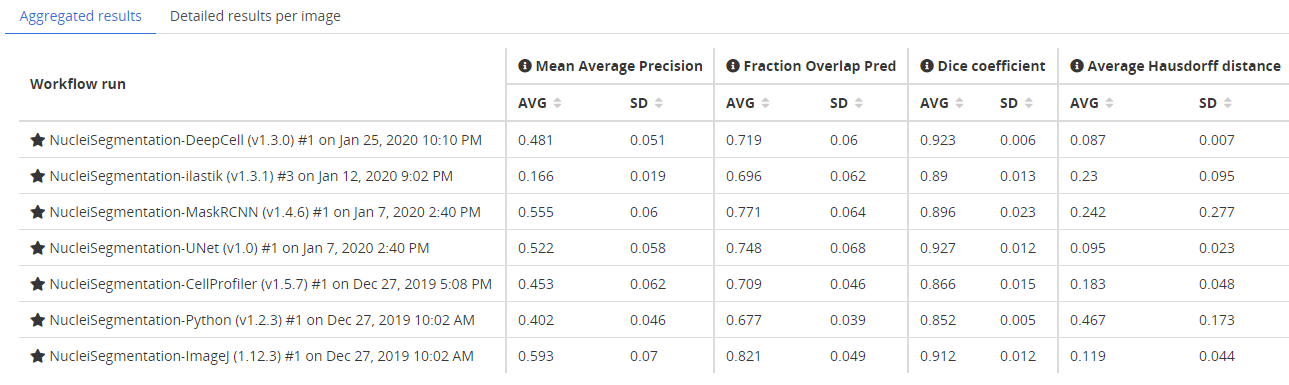

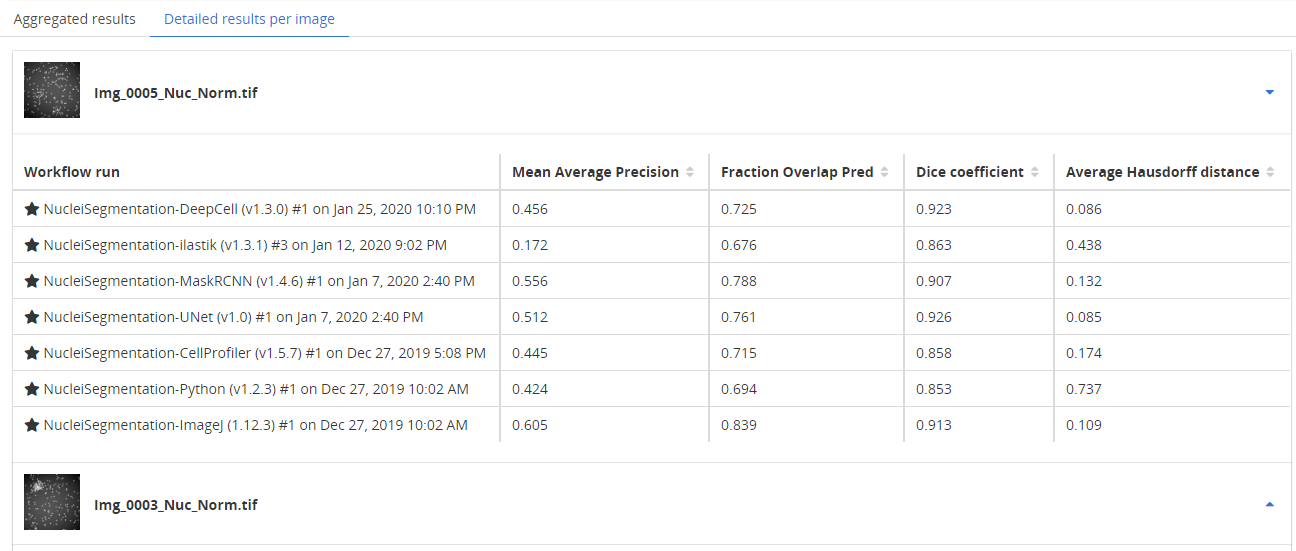

**Figure 3. BIAFLOWS workflow metric results table for the SIMCEP dataset.** Top: aggregated (all images, metrics average and standard deviation), bottom: metrics per image.

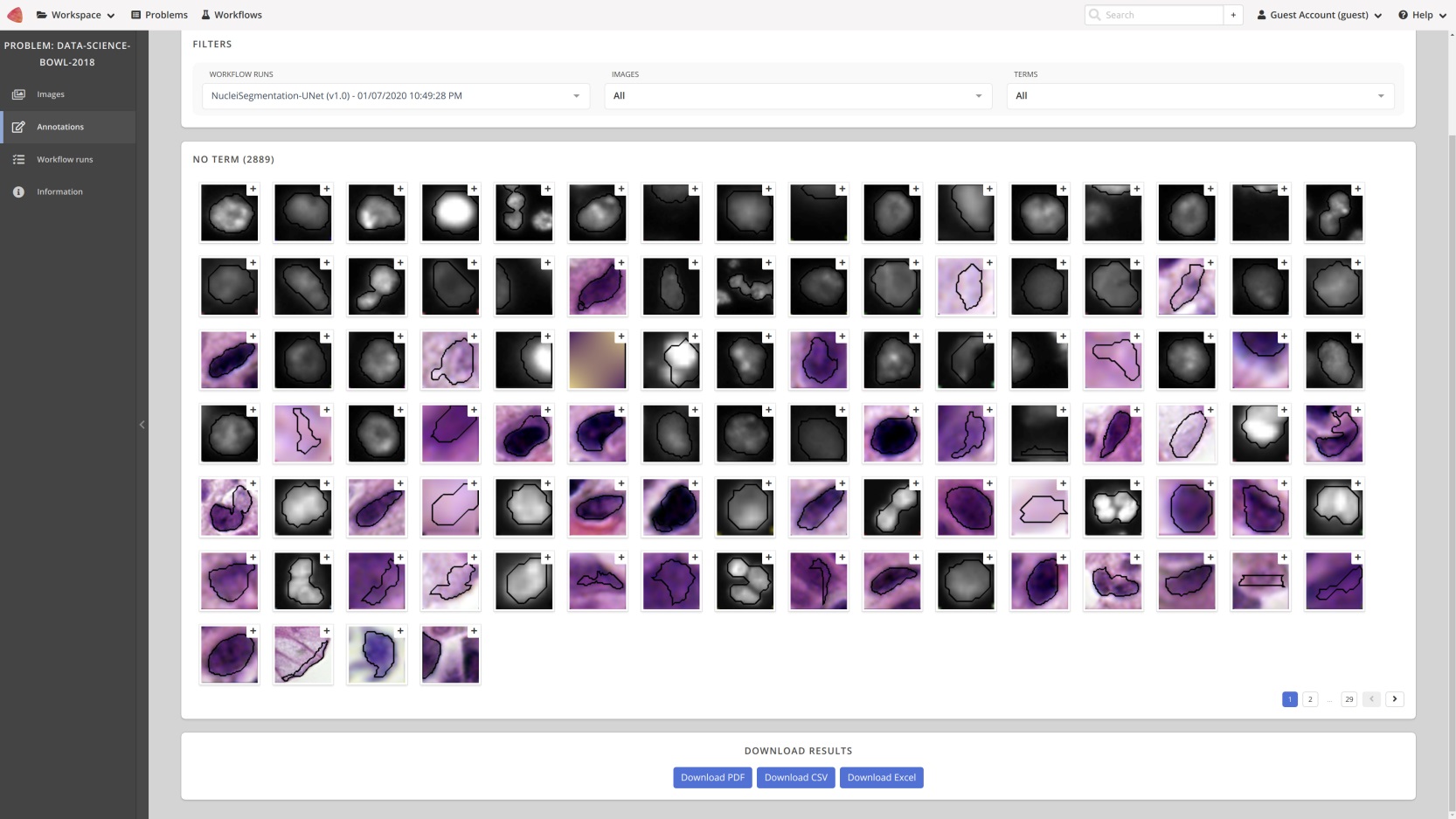

**Figure 4. BIAFLOWS gallery showing cells segmented by U-NET (first 100) from DSB dataset.** Each individual object can be clicked and visualized in context (see Fig. 5).

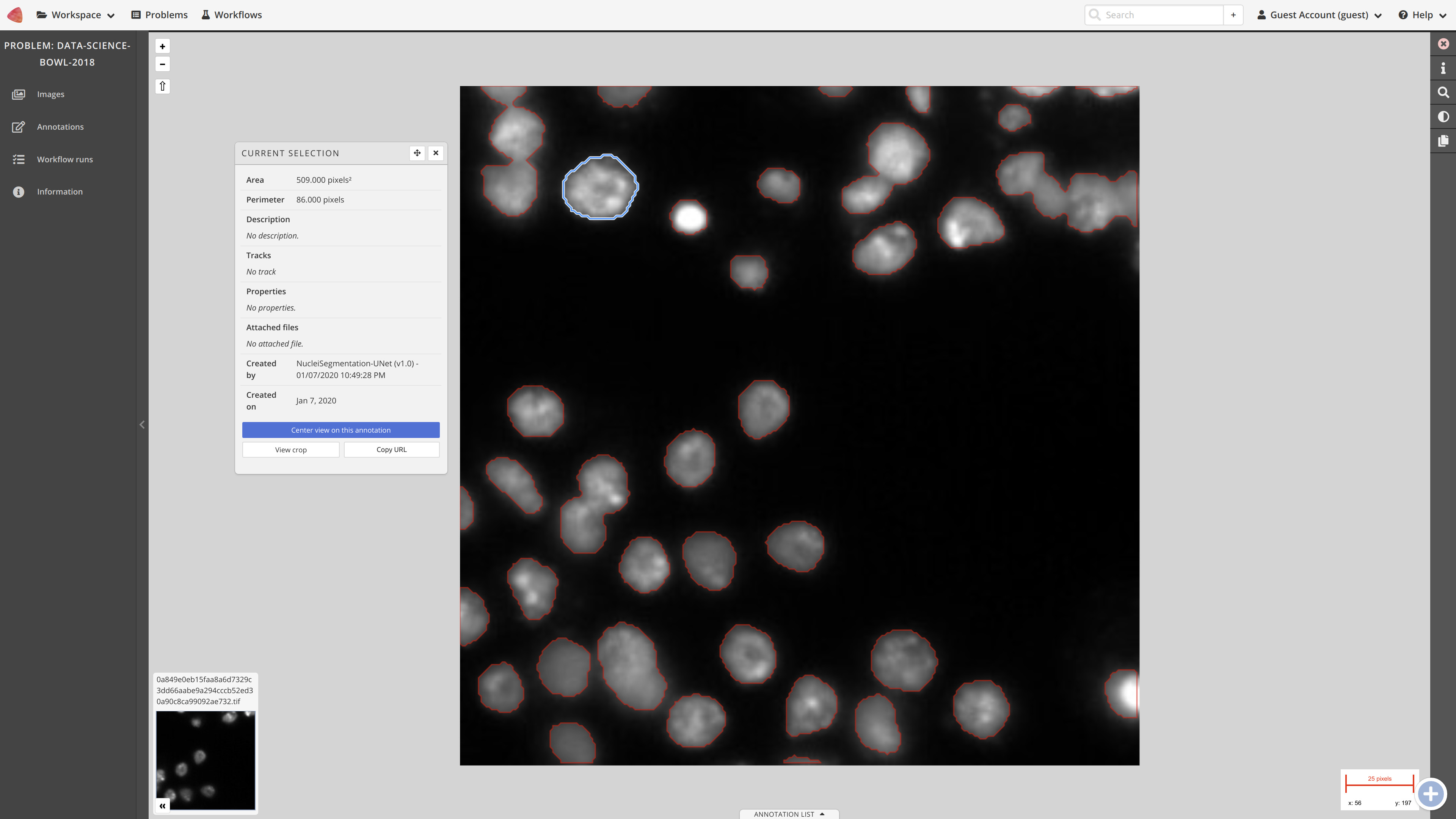

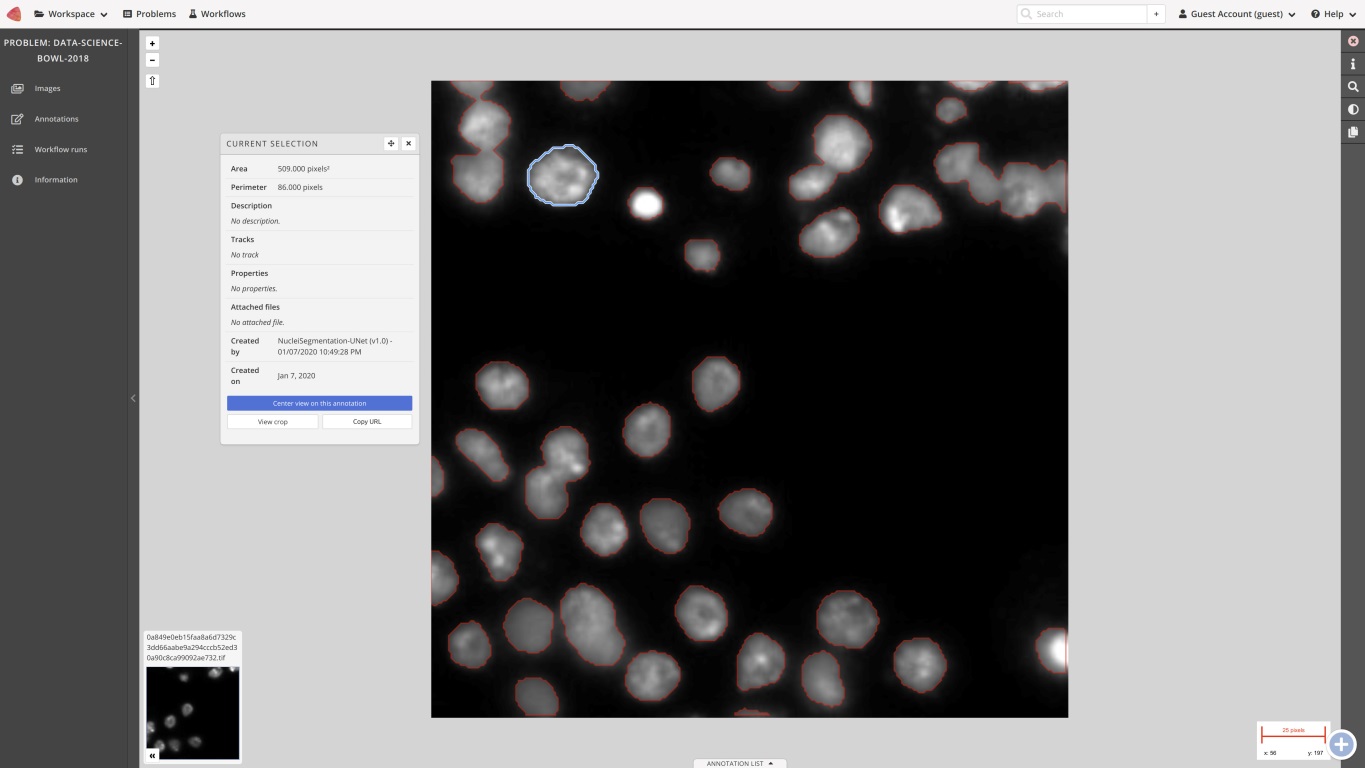

**Figure 5. BIAFLOWS image viewer displaying one image from DSB dataset and cells segmented by U-Net.** Some information on a selected cell (blue) is displayed, this cell is the first cell displayed in the gallery of Fig. 4.

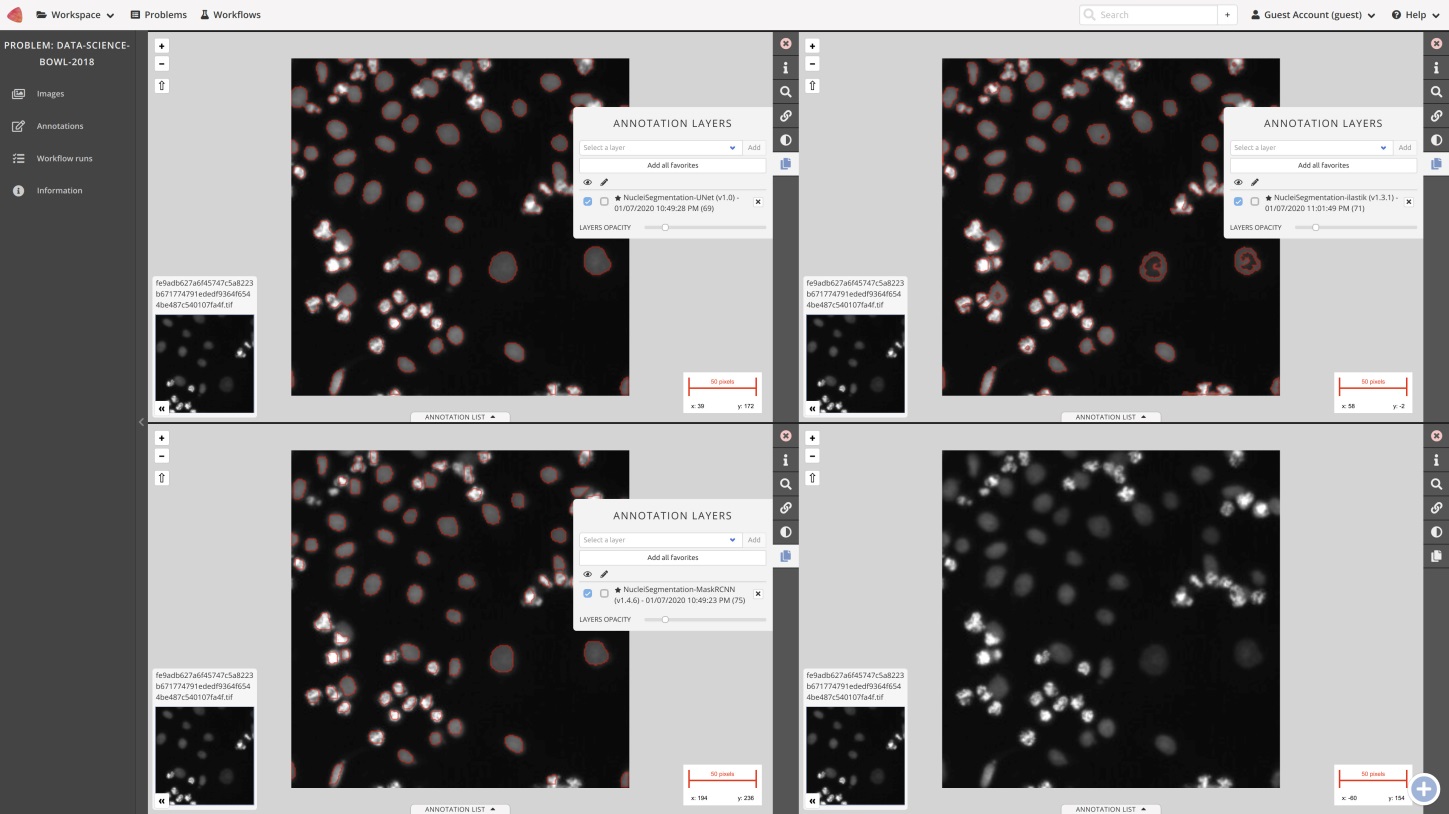

**Figure 6. BIAFLOWS image viewer side-by-side comparison of segmented cells by three workflows of one DSB image.** Top left: Unet, top right: ilastik, bottow left: MaskRCNN, bottom right: original image.

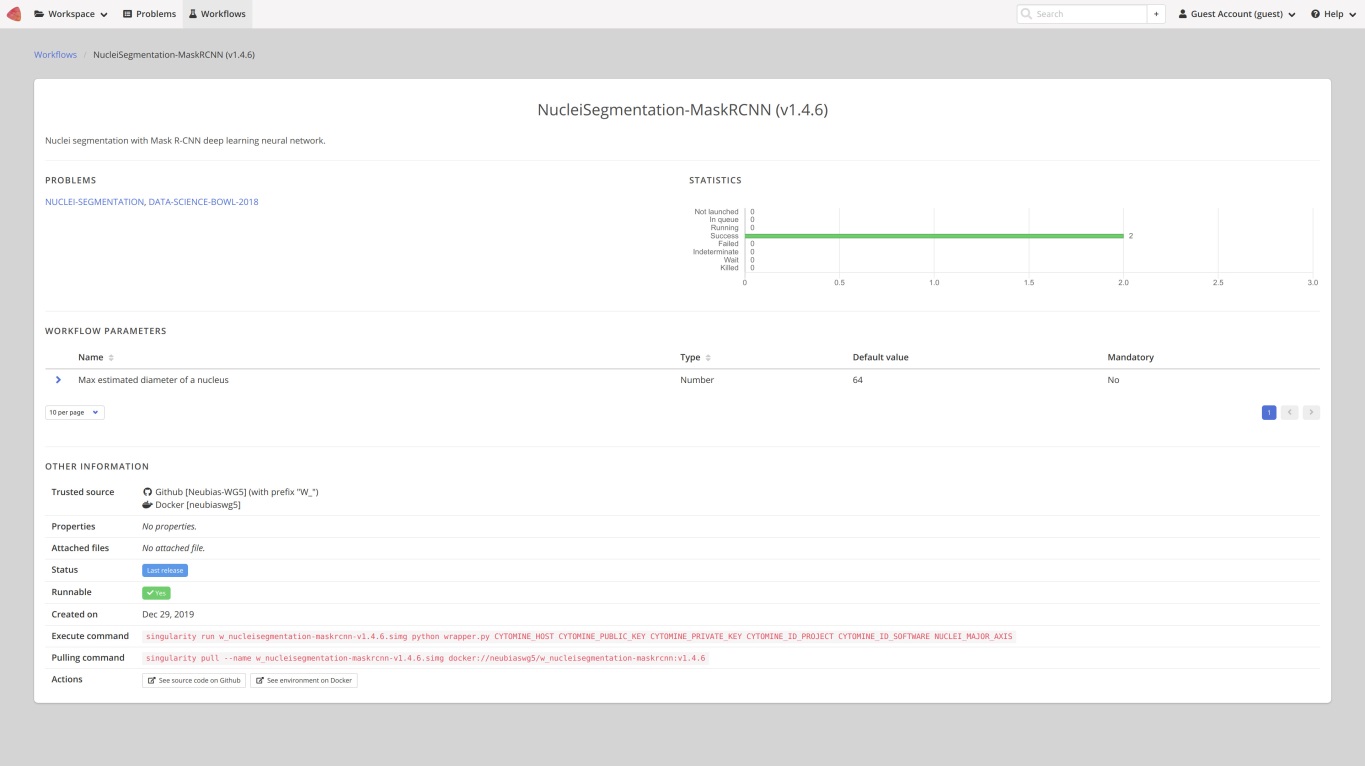

**Figure 7. BIAFLOWS workflows page.** Details of MaskRCNN workflow with direct access to original, versioned, source code on Github.

[5] A deep learning framework for nucleus segmentation using style transfer. bioRxiv 580605; doi: <https://doi.org/10.1101/580605>.

[6] <https://data.broadinstitute.org/bbbc/BBBC039/>

### Supplementary Section 3. Installing and populating BIAFLOWS locally

It is possible to install BIAFLOWS on a local server or a desktop computer. This might be useful to manage and analyse images locally or organize challenges. The procedure is described below and should take less than 30 minutes (UNIX based system recommended).

**Installing a local instance of BIAFLOWS**

The procedure described in this section is for Linux Ubuntu but it should be possible to install BIAFLOWS on other platforms (not tested). Some specific details related to deployment on Mac OS can be found [online](https://doc.uliege.cytomine.org/display/PubOp/Install+Cytomine+on+MacOS).

1/ Install requirement

BIAFLOWS runs in Docker containers, the only requirement is to install Docker.

[Check official Docker documentation to install Docker for Ubuntu.](https://docs.docker.com/install/linux/docker-ce/ubuntu/) Choose Install using the repository, set up the repository and install Docker CE.

2/ Retrieve BIAFLOWS installation files

mkdir Biaflows/
cd Biaflows/
git clone https://github.com/Neubias-WG5/Biaflows-bootstrap.git
cd Biaflows-bootstrap

3/ Configure the local instance

Edit configuration.sh and, if needed, update URLs (CORE_URL, IMS_URL, UPLOAD_URL). Make sure to use URLs that are not already used by other applications (avoid localhost) to prevent conflicts. In **/etc/hosts** of the host machine, add the following lines, adapting them accordingly to chosen XXX_URL in configuration.sh.

```
127.0.0.1   biaflows
127.0.0.1   biaflows-ims
127.0.0.1   biaflows-upload
127.0.0.1   rabbitmq
```

If needed, update data path variables (IMS_STORAGE_PATH …). All data paths must be valid and mappable in the Docker engine. If they don't exist, create all the directories (**mkdir**) corresponding to the following variables:

IMS_STORAGE_PATH
IMS_BUFFER_PATH
FAST_DATA_PATH
PROXY_CACHE_PATH
SOFTWARE_CODE_PATH
SOFTWARE_DOCKER_IMAGES_PATH
JOBS_PATH
SERVER_SSHKEYS_PATH

A reference to these URLs and paths is provided here:

<https://doc.uliege.cytomine.org/display/PubOp/Cytomine+configuration+reference>

Configure BIAFLOWS_WORKFLOWS_METRICS to *true* or *false* depending if you want to perform benchmarking: ground truth annotations are then required for all images. Setting this flag to false is the valid option if you plan to manage and process local images.

4/ Initialize the deployment

Run the installation script: sudo bash init.sh

5/ Deploy the local instance

Run the generated deployment script: sudo bash start.sh

6/ Check the running instance

When start up is finished, check that the application is running in your browser at the URL specified by CORE_URL (default: http://biaflows).

Three accounts with different access rights are automatically created (username: admin; password: admin; username: guest; password: guest; username: neubias; password: neubias). Passwords should be updated from the Account page (top right).

7/ Install sample Problems (images and ground-truth data)

After BIAFLOWS successfully installed locally, the local instance is empty. All projects available in BIAFLOWS online instance can be imported to the local instance. For this, get the public and private keys of the admin account (Account page), then run:

cd Biaflows-bootstrap

sudo bash ./inject_demo_data.sh ADMIN_PUBLIC_KEY ADMIN_PRIVATE_KEY

where ADMIN_PUBLIC_KEY and ADMIN_PRIVATE_KEY have been substituted by their respective values.

The script starts to download projects and import them in your local BIAFLOWS.

The list of imported projects can be tweaked by editing the file

Biaflows-bootstrap/configs/project_migrator/projects.txt.

The whole data injection procedure can take several minutes, depending on your Internet connection and the number of projects being imported.

**Creating a new Problem (project) in a local BIAFLOWS instance**

To create a new problem, connect as regular user or admin.

1/ Go to **Problems** tab

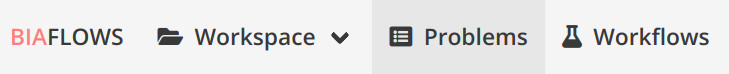

2/ Click **New Problem**

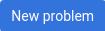

3/ Choose a meaningful problem name and save

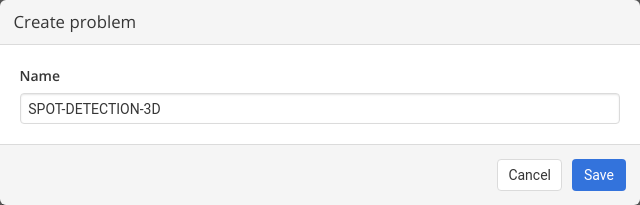

4/ The problem is ready to be configured, the following configuration is recommended

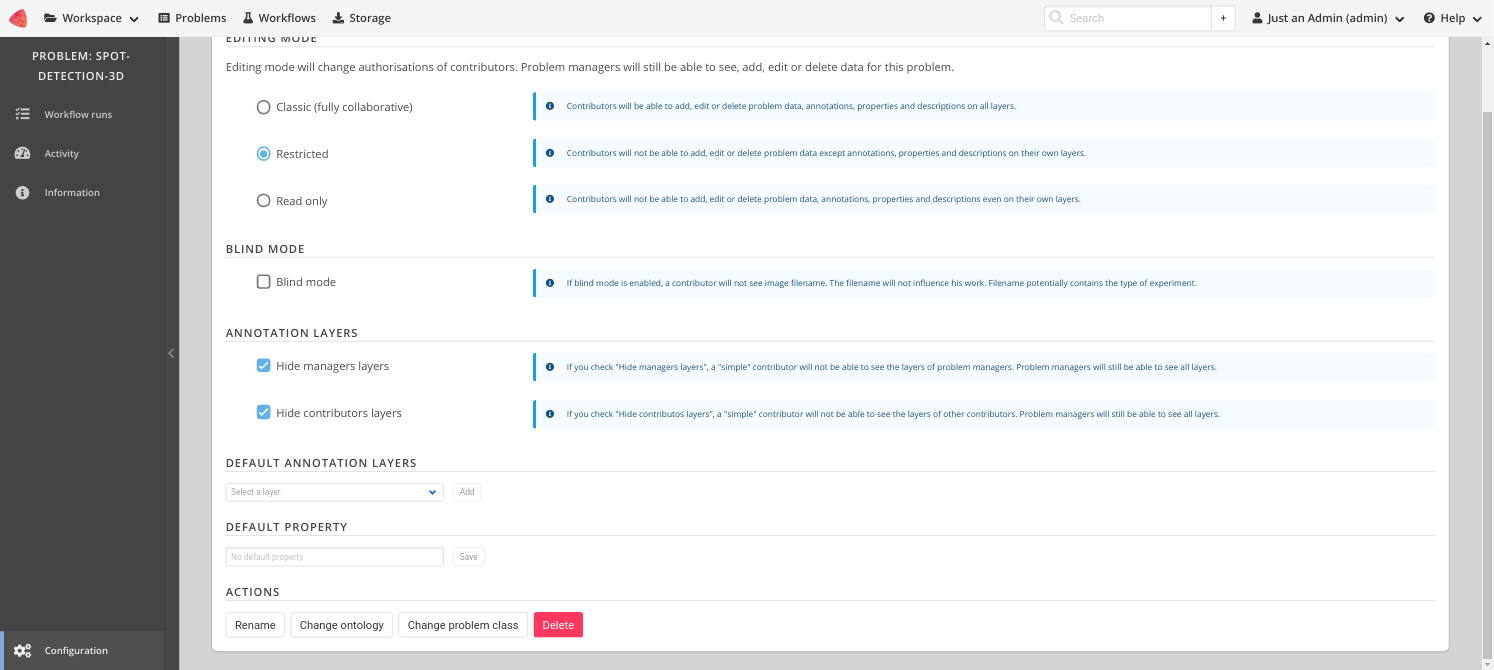

5/ Assign your problem to a problem class (see Section 5 **Problem Class,** **Ground truth annotations and reported metrics**) by clicking on **Change problem class**. The problem class specifies the format of ground truth annotations (and workflow outputs), as well as the associated benchmark metrics to be computed (if benchmark is enabled).

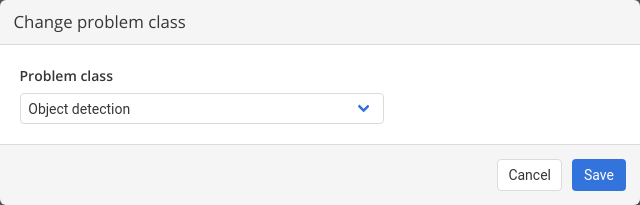

6/ Configure project members. If you work alone, you can leave contributors and project managers to default user. This can be done from the “Members” tab in the problem configuration.

7/ The problem can be fully configured to display or hide panels / tabs / tools in the user interface. This is achieved from the **Custom UI** tab in the problem configuration.

8/ A description of the problem can optionally be added from the **Information** (left sidebar). The description is displayed in **Problems** list.

**Uploading images to a local BIAFLOWS instance**

To upload new images, connect as regular user or admin.

**Supported formats**

- **2D images**: 8-bit/16-bit TIFF (or OME-TIFF files)
- **Multi-dimensional images (Z, C, T)**: single file 8-bit/16-bit OME-TIFF

**Note**: The text string **_lbl** should not be used in image names since it is a reserved field for ground truth annotation images.

1/ Go to **Storage** section

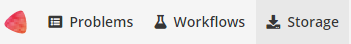

2/ Select the **Problem** to which the images should be associated with (**Link with problem**)

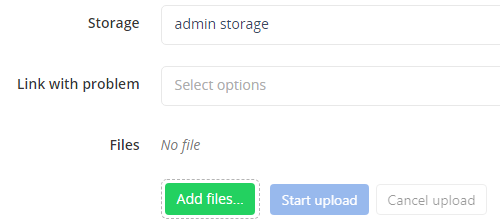

**Note**: If a problem is not in the list, make sure you are a member for this problem

3/ Click on **Add files…** and select the files from the file browser

4/ Start upload with **Start upload** and wait until completion

The status can be:

- **DEPLOYED/CONVERTED**: The image is correctly imported to BIAFLOWS
- **ERROR FORMAT**: The file format is not supported
- **ERROR EXTRACTION**: Something went wrong during metadata extraction
- **ERROR CONVERSION**: Something went wrong during the conversion of the image into the BIAFLOWS internal image format
- **ERROR DEPLOYMENT**: Something went wrong during the communication with BIAFLOWS API. It can be due to access rights, or other unexpected error

**Note**: Images uploaded to storage can also be associated to a Problem after upload (Problem: **Add image**). This can be useful to associate the same image to several Problems.

**Uploading ground truth annotations to an existing BIAFLOWS problem**

If you plan to perform benchmarking, ground truth annotations should also be uploaded and associated to every image of a problem. The format of these annotations depends on the associated problem class (see Section 5 **Problem Class**, g**round truth annotations and reported metrics**).

Image annotations (e.g. binary masks) should be uploaded as 16-bit TIFF (or OME-TIFF) for 2D images and as single file 16-bit OME-TIFF for multidimensional (C,Z,T) images. They should be uploaded by following the procedure described in the previous section and by setting the same name as their corresponding image + **_lbl** suffix (e.g. **AnImage.ome.tif** and **AnImage_lbl.ome.tif**).

Other required annotations (e.g. SWC, division text file) should be added to the images as attached files. To do so, expand the image (blue arrow) in the list and click on **Add** next to **Attached files**. These can also be added programmatically using our Python client.

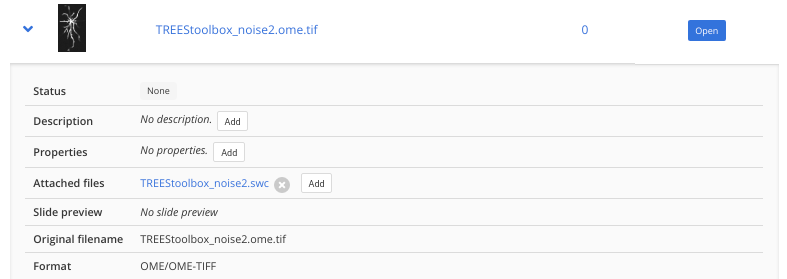

**Adding existing workflows from trusted sources to a local BIAFLOWS instance**

It is possible to integrate existing BIAFLOWS workflows to any BIAFLOWS instance. This operation requires configuring an external trusted source made of:

1. A source code registry (typically a Github user space)
2. An execution environment registry (typically a DockerHub user space)

If your workflow repositories are mixed with other repositories in your user space, you can specify a prefix to distinguish workflows repositories. For instance, all bioimage analysis workflows developed by NEUBIAS are prefixed by **W_** and available from this user space: [https://github.com/Neubias-WG5](https://github.com/neubias-wg5).

Some information regarding trusted sources is given below.

To manage trusted sources, you need to be administrator.

1/ Connect as administrator by clicking Open admin session:

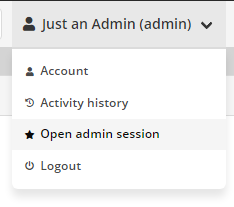

2/ In administration page, go to **Trusted sources** tab and click **Add trusted source**

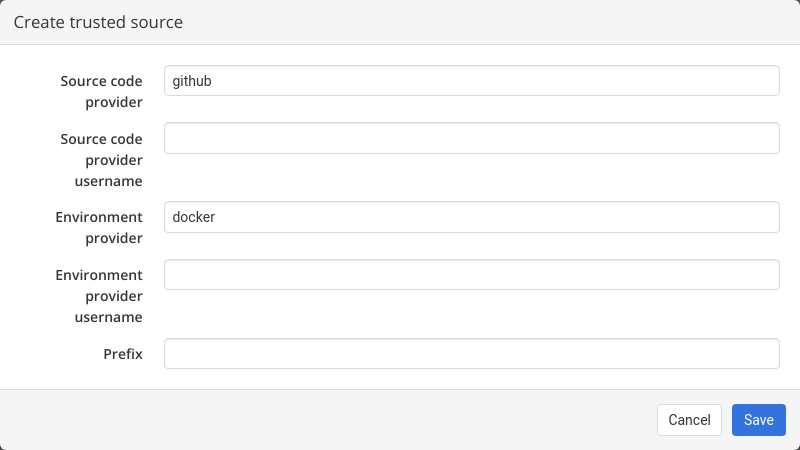

3/ Fill the form and **Save**

For instance, to add NEUBIAS curated set of workflows, the trusted source has to be configured as follows:

- **Source code provider:** github
- **Source code provider username:** Neubias-WG5
- **Environment provider:** docker
- **Environment provider username:** neubiaswg5
- **Prefix:** W_

4/ Trusted sources are periodically checked (about every 5/10 minutes) to automatically add new versions of existing workflows or new workflows, but you can also click on **Refresh** to trigger the check.

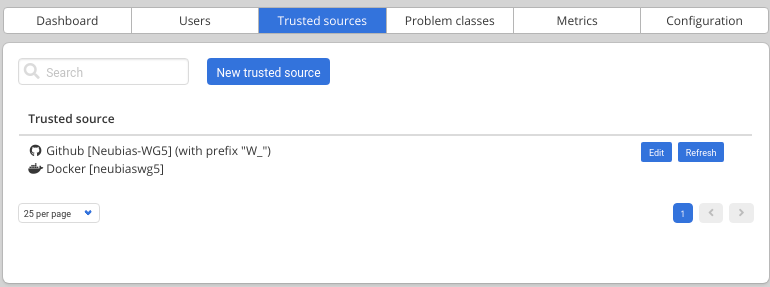

5) Once a workflow is imported, it has to be linked to a BIAFLOWS **Problem**. This can be performed in the Configuration panel of the **Problem** (**Workflows** tab) by toggling **Enable** for that workflow as illustrated below:

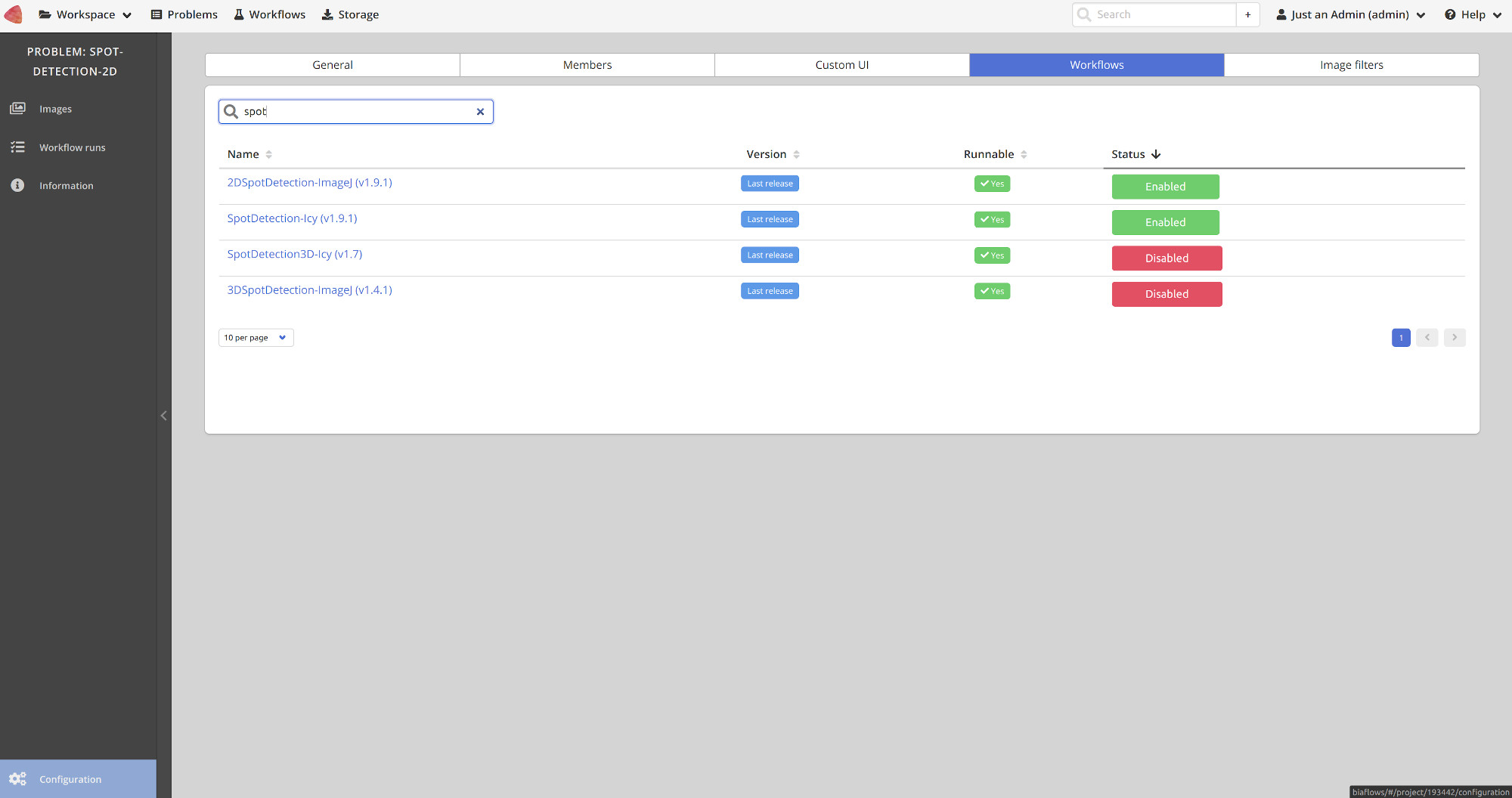

### Supplementary Section 4. Creating a BIA workflow and adding it to a BIAFLOWS instance

**Introduction**

BIAFLOWS workflows are *Docker images* encapsulating a complete execution environment together with a workflow addressing a BIA Problem. These *Docker images* can be compiled automatically online. BIAFLOWS instances automatically fetch new workflows and make them available from the user interface. Sample workflows running in ImageJ (macros and scripts), ICY, CellProfiler, ilastik, Vaa3D, Python, Octave and Jupyter notebooks can be found in this GitHub repository: <https://github.com/neubias-wg5>. The procedure to package a workflow and add it to a BIAFLOWS instance is described in this section. Users willing to get help can write to [https://forum.image.sc](https://forum.image.sc/) forum or contact.

**BIA workflow requirements**

BIAFLOWS workflows must:

- Run headless from command line
- Take an input folder of 8 bit/16 bit TIFF (2D) or single file OME-TIFF (C,Z,T) images
- Expose functional parameters and parse them from command line call
- Export results to an output folder in a format specified for the Problem Class (see **Problem Class, ground truth annotations and reported metrics**).

The workflow and its software execution environment are fully defined from a set of 4 files:

●	A Dockerfile configuring software execution environment (OS, libraries, software...)

●	The workflow executable or, more commonly, a script running on a BIA platform

●	A Python script (wrapper.py), sequencing operations (*Docker image* entry point)

●	A descriptor (descriptor.json) specifying workflow parameters and default values.

**Step 1.** Create a workflow GitHub repository

Create a workflow repository in a GitHub source trusted by the BIAFLOWS instance you plan to add the workflow to. The names of workflow repositories should start by a fixed prefix (**W_** recommended since it is the convention used by BIAFLOWS online instance) and hold no space.

**Step 2.** Add the 4 required files to the workflow repository

It is recommended to reuse existing files from similar workflow repositories in <https://github.com/Neubias-WG5>. For this, follow these guidelines:

- A descriptor from the **Problem Class** you target (e.g. Object Segmentation)
- A DockerFile configuring the BIA platform you target (e.g ImageJ)
- A wrapper script from the **Problem Class** and the **workflow type** you target.

Note: The flag **is_2d** should be used to specify if the images are strictly 2d or multidimensional.

The following workflow types have already been tested and are available from <https://github.com/Neubias-WG5>: ImageJ / FIJI macro, ImageJ Python script, ICY protocol, CellProfiler pipeline, Octave script, ilastik pipeline, Vaa3D plugin, Python 2.X or 3.X script based on Scikit-learn or KEras/Pytorch.

**Step 3.** Update the following sections of the **Descriptor**

**Workflow and associated Docker image names**

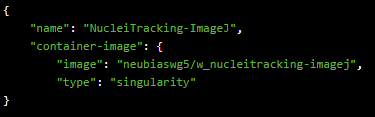

Update *name* to match GitHub workflow repository name (without prefix)

Update *image* to match the name of your workflow GitHub repository (lower case only)

**Command line call of the Docker image**

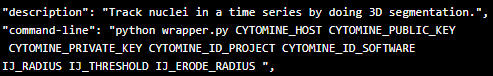

*Description*: Update workflow description

*Command-line*: Update parameter list (here last 3 arguments)

**Workflow parameter sections**

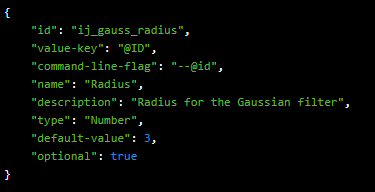

Update / add as many parameter sections as required to match the parameter list from command line call.

*id*: should match parameter name in command line call (lower case)

*name*: name that will appear in BIAFLOWS user interface (parameter dialog box)

*description*: context help in BIAFLOWS user interface (parameter dialog box)

*type*: String or Number

*default-value*: the default value in BIAFLOWS user interface (parameter dialog box).

**Step 4.** Update DockerFile

Update the line copying the workflow from the GitHub repository to the workflow Docker image, for instance:

ADD NucleiTracking.ijm /fiji/macros/macro.ijm

If necessary, append commands to install additional required libraries/plugins to the execution environment.

**Step 5.** Update wrapper script

Update workflow command line call in wrapper.py.

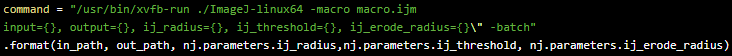

Update/add parameters to match parameters defined in JSON descriptor (Step 2).

**Step 6.** Adapt your workflow script

Adapt your workflow script to fulfil workflow requirements and parse parameters from command line. For instance for an ImageJ macro:

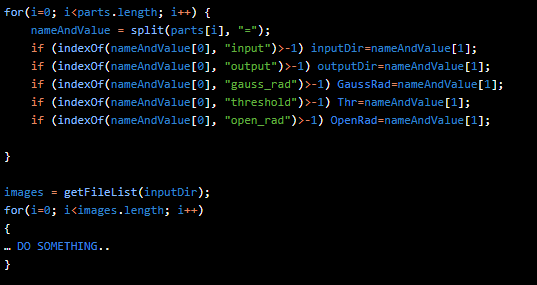

**Step 7.** Create Docker image in DockerHub

Sign in to DockerHub and create a new public repository. The repository name must match the container-image name used in Step 3.

**Step 8.** Link repository to workflow GitHub repository and configure workflow *Docker image* automated build according to the following example:

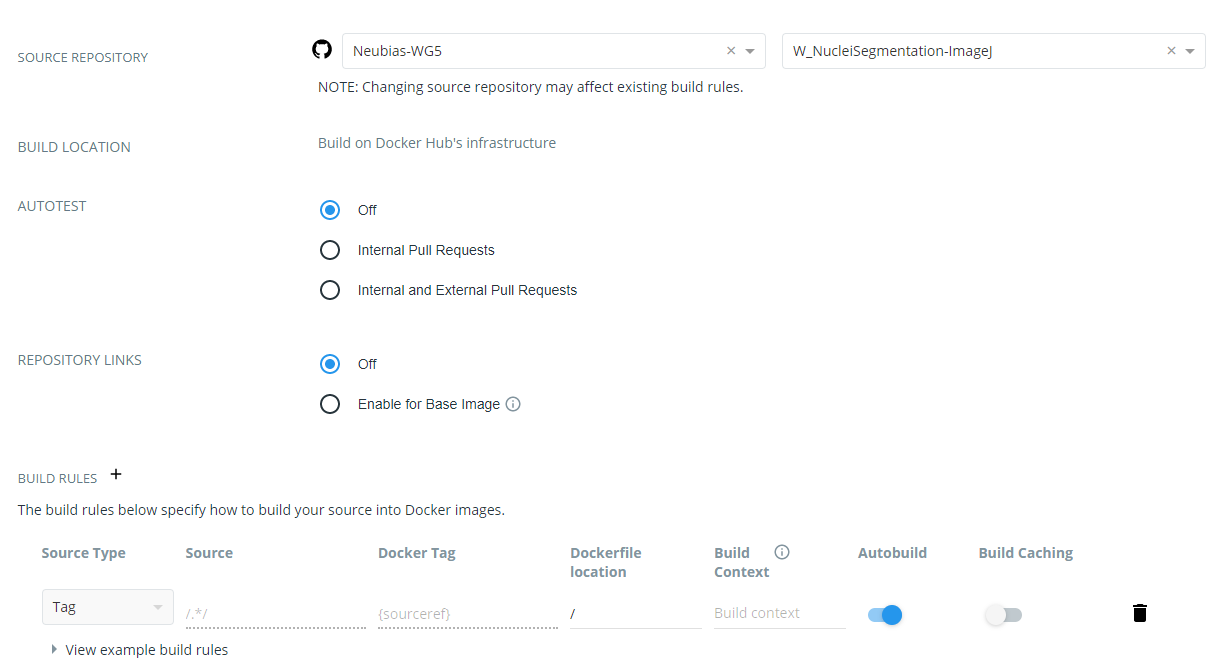

**Step 9.** Trigger a workflow release

Trigger a release from GitHub workflow repository with version tag such as 0.1, 0.2, 1.0...

**Step 10.** Workflow Docker image build

Check from DockerHub that the workflow *Docker image* has built successfully. If not, parse the log and fix issues by modifying DockerFile and retriggering a new release.

**Step 11.** Add workflow to BIAFLOWS problem

Once the *Docker image* is built, a BIAFLOWS instance fetches the image from the trusted source and make it available (possibly after up to 5/10 minutes). Sign in as administrator to BIAFLOWS and browse to the **Problem** you want to add the workflow to. Then, click on the **Configuration** icon (bottom left of the side bar).

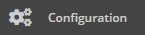

Search for the workflow (recently added workflows are on top of the list) and enable it. Older workflow versions can be disabled if this is an update to an existing workflow.

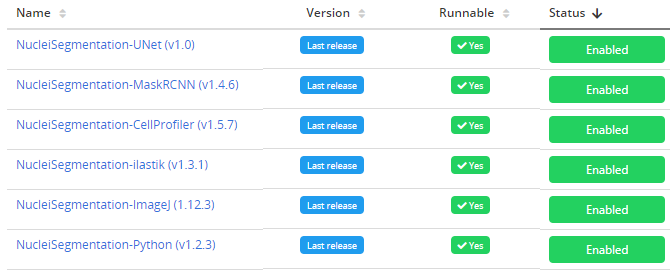

**Step 12.** Run the workflow

Test the workflow by running it from BIAFLOWS / **Workflow runs** (requires execution rights).

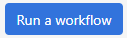

 If execution fails, read the execution log, update the code and trigger a new release.

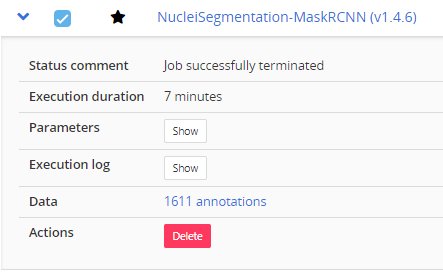

**Detailed Developer guide**

This section provides some more details on BIAFLOWS workflows and details how to compile and debug BIAFLOWS workflows Docker image locally and add them to an existing BIAFLOWS instance.

**Details on Python wrapper script and JSON descriptor**

The sequence of operations commonly performed by BIAFLOWS Python wrapper scripts is detailed in Table S4.1. All workflows provided in BIAFLOWS repository follow this template. A complete reference to BIAFLOWS workflows JSON descriptor can be found [online](https://doc.uliege.cytomine.org/display/ALGODOC/Software+JSON+descriptor+reference)[.](https://doc.cytomine.be/display/ALGODOC/Software+JSON+descriptor+reference)

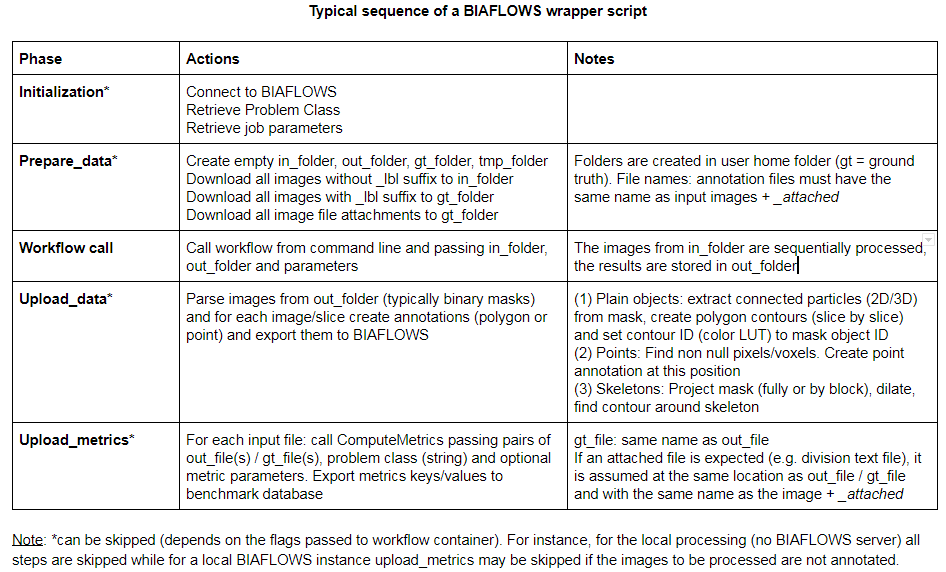

**Figure S4.1. Typical steps of a BIAFLOWS Python wrapper script**

**Installing software required for development (only once)**

As workflows run inside a Docker container and since their Python wrapper script interacts with a BIAFLOWS instance, it is required to install Docker and Python 3 on your local machine. Our Python client is also required for development.

**Docker installation instructions can be found here:**

For Linux:

<https://www.digitalocean.com/community/tutorials/how-to-install-and-use-docker-on-ubuntu-18-04>

For Windows:

<https://docs.docker.com/docker-for-windows/install/#install-docker-for-windows-desktop-app>

**Python 3 and Cytomine Python client instructions can be found here:**

See <https://doc.uliege.cytomine.org/display/ALGODOC/Data+access+using+Python+client>

In the following steps, we will use the workflow “NucleiSegmentation-ImageJ” as reference:

<https://github.com/Neubias-WG5/W_NucleiSegmentation-ImageJ>

**Step 1**. **Uploading a new workflow descriptor to BIAFLOWS**

Workflows have first to be described through a JSON descriptor, e.g.:

<https://github.com/Neubias-WG5/W_NucleiSegmentation-ImageJ/blob/master/descriptor.json>

Currently, some sections have to be customized manually, and some conventions must be respected to allow automatic parsing by BIAFLOWS. We recommend using <https://github.com/Neubias-WG5/W_Template/blob/master/descriptor.json> as template for your JSON descriptor.

Choose a workflow name without space. The description field (supporting restricted HTML) should be filled to document the workflow and it will be displayed from BIAFLOWS UI.

As inputs (workflow parameters), the five parameters (cytomine_host, cytomine_public_key, cytomine_private_key, cytomine_id_project, cytomine_id_software) are mandatory.

The workflow parameters should also be described:

- id: the parameter name (e.g : “ij_radius”)
- value-key: a reference for the parameter in the command line. Keep “@ID”, which is a shorthand meaning “replace by the parameter id, in uppercase”. In our example, it will be replaced at parsing time by “IJ_RADIUS”
- command-line-flag: At execution time, the value-key in the command line will be replaced by the command-line-flag followed by the parameter value. Keep “--@id”. In our example, it will be replaced in the command line by “--ij_radius”.
- name: a human readable name displayed in BIAFLOWS
- type: Number, String, Boolean
- optional: set to true only if the workflow execution is not influenced by the presence or the absence of the parameter (e.g a “verbose” parameter). Workflow parameters having an influence on the results should never be optional.
- default-value: the default value of the parameter (in BIAFLOWS interface).

Do not forget to update the command line, with the parameter value keys. For instance, for workflow parameters ij_radius and ij_threshold:

python wrapper.py CYTOMINE_HOST CYTOMINE_PUBLIC_KEY CYTOMINE_PRIVATE_KEY CYTOMINE_ID_PROJECT CYTOMINE_ID_SOFTWARE IJ_RADIUS IJ_THRESHOLD

To make a workflow available from a BIAFLOWS instance, it is currently required to publish its descriptor using Cytomine Python client. This can be performed by running the following Python code inside the folder holding the JSON descriptor you have created:

```
from cytomine import Cytomine
from cytomine.utilities.descriptor_reader import read_descriptor
with Cytomine(host, public_key, private_key) as c:
	read_descriptor("descriptor.json")
```

***host*** is the url of your BIAFLOWS server, e.g. [https://biaflows.neubias.org](https://biaflows.neubias.org/)

***public_key*** and ***private_key*** can be found from user Account page (section API KEYS)

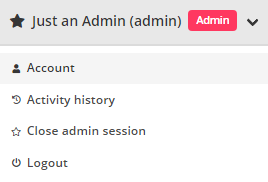

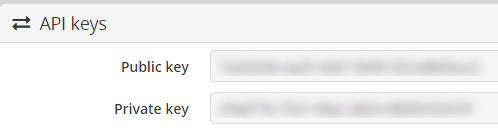

**Step 2. Linking a new workflow to a BIAFLOWS project**

- From *Problems*, select the problem to which you want to add the workflow
- Go to *Problems* > *Configuration* > *Workflows* and enable the workflow

For now, as the workflow has been added manually, it will be referenced as **Not Runnable** and no version information will be provided from the UI.

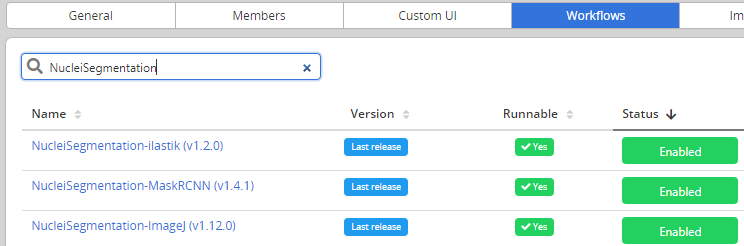

Next, Go to *Projects* > *Configuration* and make sure that Jobs tab is activated (green)

**Step 3.** **Creating the Dockerfile**

Docker files specify the execution environment. They typically start by creating (FROM) a layer from an existing Docker image with basic operating system. Then they execute commands (RUN) to install specific software and libraries, and copy (ADD) files (e.g. the Python wrapper script and workflow script) into the execution environment the workflow will be called from. Finally, the ENTRYPOINT is set to the wrapper script.

A sample DockerFile is available here:

<https://github.com/Neubias-WG5/W_NucleiSegmentation-ImageJ/blob/master/Dockerfile>

If you do not know how to configure the Dockerfile, it is recommended to adapt the Dockerfile from an existing BIAFLOWS workflow using the same target software (e.g. an ImageJ macro).

**Note**: If you create a Dockerfile from scratch, always use the most accurate tag when referring to an existing Docker image (e.g. prefer python:3.6.9-stretch over python:3.6). If the tag is not accurate, the underlying docker image could change over time, heavily impairing reproducibility!

**Step 4. Creating the wrapper script**

It is recommended to adapt a wrapper script: 1) from same problem class, 2) processing image of same dimensionality (e.g. 3D), and 3) matching the software you are planning to use (e.g. ImageJ macro). In this case, only the workflow call (command line) needs to be adapted. A sample wrapper script is available here:

<https://github.com/Neubias-WG5/W_NucleiSegmentation-ImageJ/blob/master/wrapper.py>

Note: The flag **is_2d** should be used to specify if the images are strictly 2d or multidimensional.

**Step 5. Building the workflow image, running it in a local container and debugging**

A new workflow can be directly pushed to GitHub and be built in DockerHub, but it is preferable to test it locally beforehand. For this, it is required to build and run the Docker image locally:

Building the container (you need at least around 5GB disk space for this operation)

From a directory where you gathered the 4 files required to describe the workflow:

cd ~/Documents/Code/NEUBIAS/W_NucleiSegmentation-ImageJ$

sudo docker build -t seg2d .

Here seg2d is the name of the Docker image to build locally.

Running the Docker image:

sudo docker run -it seg2d --host *host* --public_key *public_key* --private_key *private_key* --software_id *software_id* --project_id *project_id* --ij_threshold 15 --ij_radius 4

The list of command-line parameters should exactly match the parameters defined in the JSON descriptor file. BIAFLOWS instance URL and credentials should also be filled, as well as valid **workflow_id** (using **--software_id**) and **problem_id** (using **--projet_id**).

These IDs can be retrieved from the URL bar while respectively clicking on a problem (from BIAFLOWS **Problems** tab) and on a workflow (from BIAFLOWS **Workflows** tab):

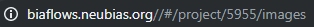

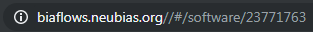

In this example, workflow_id=23771763 and problem_id=5955.

If a workflow fails at execution this is reported in **Workflow runs** section. Some **Execution log** can be downloaded by expanding a workflow run from the blue arrow:

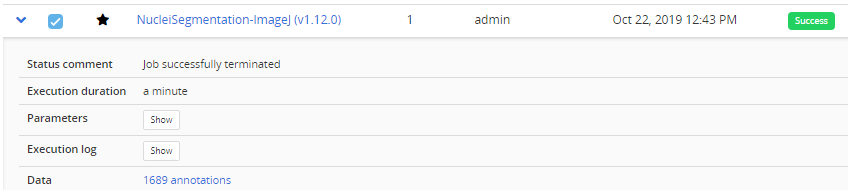

In that case, no associated benchmark metric is associated to this run. There is hence no risk that this would be left unnoticed by the user. For debugging, Docker can be run with an interactive session:

sudo docker run --entrypoint bash -it seg2d

If needed, it is also possible to launch the Docker with X enabled, e.g. to debug imageJ macro more easily:

xhost + sudo docker run --entrypoint bash -v

/home/yourusername/tmp/test:/data -e DISPLAY=$DISPLAY -v /tmp/.X11-unix:/tmp/.X11-unix -it seg2d

If you want to access local images without having to download them each time from BIAFLOWS, you can also attach a local folder to a folder inside the Docker container (-v option), for instance:

sudo docker run --entrypoint bash -v /home/yourusername/tmp/test:/data -it seg2d

Some other useful Docker commands

Check if an image is running:	ps -a

Kill a running container:	sudo docker rm 65e88b2015df

Kill all running containers:	sudo docker rm $(sudo docker ps -a -q)

Download a specific container	sudo docker pull neubiaswg5/fiji-base:latest

Note: To download a recently updated workflow image, it is necessary to first remove older versions manually.

**Step 6. Publishing a workflow with version control**

Once your workflow is running properly, you can officially publish it with version control.

To allow automatic import to BIAFLOWS, the set of files previously described should be stored in a GitHub repository (linked to DockerHub) from an account trusted by the target BIAFLOWS instance.

The Github repository name **must be given by:**

Github repo name = {prefix}{workflow_name}

where

- {prefix} is an optional prefix for the trusted source (see **Installing and populating BIAFLOWS locally**)
- {workflow_name} is the name of the workflow as given in the “name” field in the JSON descriptor (see Step 1).

For instance, for a trusted source with a prefix **W_**: **W_NucleiSegmentation-ImageJ**.

Adding/editing trusted sources is performed from Admin / Trusted sources (**Installing and populating BIAFLOWS locally**):

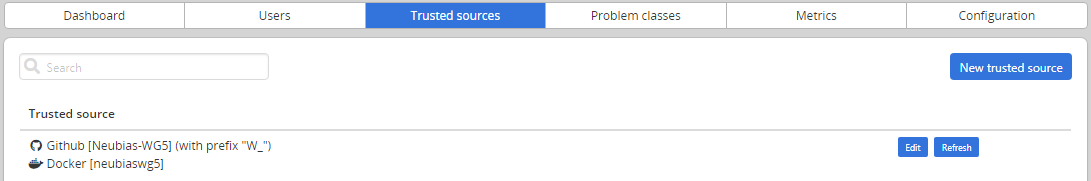

**Step 7**. **Linking a GitHub repository to DockerHub (only once)**

We assume that you created a trusted GitHub organization (e.g. **neubias-wg5**) and a workflow repository holding the 4 workflows files. It is now required to link DockerHub to GitHub. Fortunately, this operation has to be performed only once for a given GitHub organization:

1. Create an account on DockerHub : <https://hub.docker.com/> and login
2. Create an automatic build by linking Docker account to GitHub organization account
3. In DockerHub website, click on Create > Create Automated Build

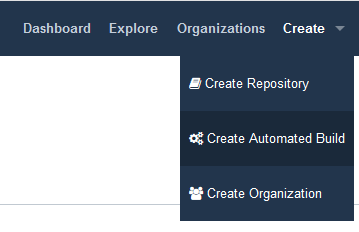

4. In **Linked Accounts,** click on **Link Github**

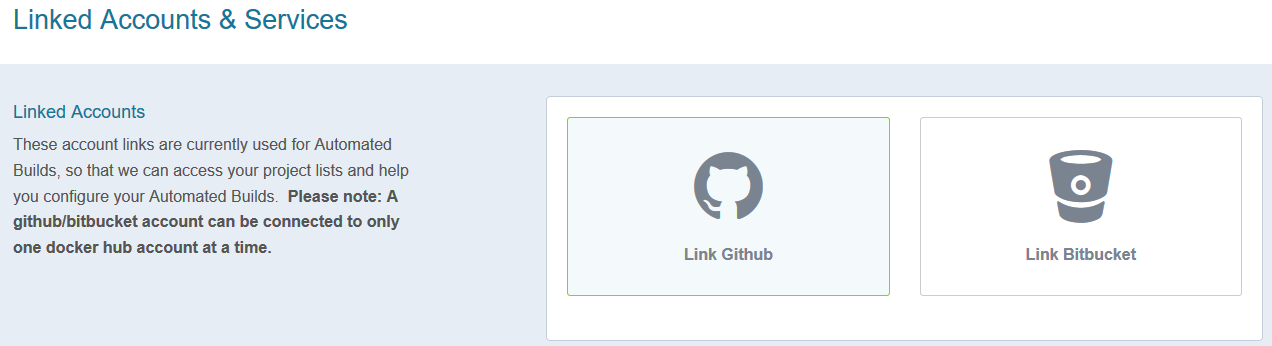

5. Click **Select**
6. Ensure that Organization access (e.g. **Neubias-WG5)** is selected (green check mark) and click on **Authorize docker**
7. Enter your GitHub password to enable access

**Step 8**. **Associating a new workflow repository to DockerHub**

Once your Github organization account and DockerHub are linked, it is possible to create an automated build procedure for each workflow. This procedure will build a workflow Docker image each time a new release is triggered from a GitHub workflow repository. This image is automatically downloaded by the BIAFLOWS instance and the new workflow version will be available for the target problem.

To do so, from DockerHub:

1. Click on Create > Create Repository+
2. In build settings click on GitHub icon
3. Select organization (e.g. **neubiaswg5**) and workflow Github repository (e.g. **W_NucleiSegmentation-ImageJ**) at the bottom of the page
4. Choose the Docker registry repository name. In practice, keep the same as Github repository (DockerHub will convert uppercase letters into lowercase).
5. Enter a short description (less than 100 characters) and click **Create**
6. Click on *Click here to customize the build settings* and configure as in figure below
7. Click on Save

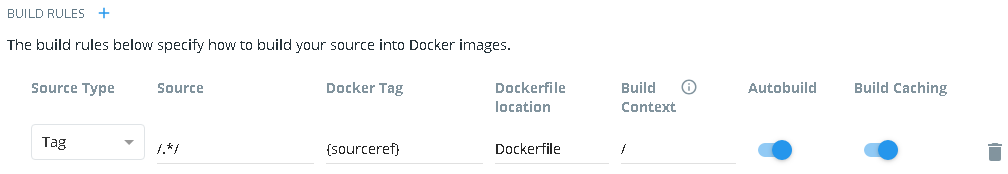

The DockerHub repository name must be reflected in the JSON descriptor:

image = {dockerhub_organization}/{github_repo_name.toLowerCase()}

For example, in the JSON descriptor:

**container-image**:

{

- **image**: "neubiaswg5/w_nucleisegmentation-imagej",
- **type**: "singularity"

},

**Step 9. Creating a versioned release on GitHub**

To create versioned releases of the workflow, go to GitHub and draft a new release (see <https://goo.gl/bFz66N>). This will add a new tag to the last commit. As we configured automatic build in previous step, a new Docker image will be built and published with the same tag. BIAFLOWS instances trusting this GitHub / DockerHub repository will now automatically fetch and make this new version available from the UI (possibly after up to 5/10 minutes).

### Supplementary Section 5. Problem class, ground truth annotations and reported metrics

To perform benchmarking, ground truth annotations should be encoded in a format that is specific to the associated problem class. BIA workflows are also expected to output results in the same format. Currently 9 problem classes are supported in BIAFLOWS and their respective annotation formats and computed benchmark metrics are described below.

**Note**: each problem class has a long name (explicit) and short name, for instance Object Segmentation (ObjSeg). The same hold for metrics, for instance DICE (DC).

See section **Benchmarking Metrics** for metrics description.

**Problem class: Object Segmentation (ObjSeg)**

Task: Delineate objects or isolated regions

Object Encoding: 2D/3D label masks with foreground > 0 (one unique ID per object), background = 0

Reported metrics: DICE (DC), AVERAGE_HAUSDORFF_DISTANCE (AHD), computed by [VISCERAL](http://www.visceral.eu/resources/evaluatesegmentation-software/) executable (archived [here](https://github.com/Neubias-WG5/neubiaswg5-utilities/blob/master/bin/Visceral)), Fraction overlap (FOVL) computed by [custom Python code](https://github.com/Neubias-WG5/neubiaswg5-utilities), Mean Average Precision computed by [Data Science Bowl 2018](https://www.kaggle.com/c/data-science-bowl-2018/overview/evaluation) Python code.

**Problem class: Spot / object counting (SptCnt)**

Task: Estimate the number of objects

Object Encoding: 2D/3D binary masks, exactly 1 spot/object per non null pixel

Reported metrics: RELATIVE_ERROR_COUNT (REC), computed by [custom Python code](https://github.com/Neubias-WG5/neubiaswg5-utilities).

**Problem class: Spot / object detection (ObjDet)**

Task: Detect objects in an image (e.g. nucleus)

Object Encoding: 2D/3D binary masks, exactly 1 object per non null pixel

Reported metrics: CONFUSION_MATRIX (TP, FN, FP), F1_SCORE (F1), PRECISION (PR), RECALL (RE), Distance RMSE (RMSE), computed by [Particle Tracking Challenge](http://bioimageanalysis.org/track/) metric Java code (particle matching only, [archived here](https://github.com/Neubias-WG5/neubiaswg5-utilities) in bin / DetectionPerformance.jar)

**Problem class: Pixel/Voxel Classification (PixCla)**

Task: Estimate pixels class

Object Encoding: 2D/3D class masks, gray level encodes pixel/voxel class, background = 0

Reported metrics: F1_SCORE (F1), ACCURACY (ACC), PRECISION (PR), RECALL (RE), computed by [custom Python code](https://github.com/Neubias-WG5/neubiaswg5-utilities)

**Problem class: Filament Tree Tracing (TreTrc)**

Task: Estimate the medial axis of a connected filament tree network (one per image)

Object Encoding: [SWC file](http://www.neuronland.org/NLMorphologyConverter/MorphologyFormats/SWC/Spec.html)

Reported metrics:

UNMATCHED_VOXEL_RATE (UVR), computed by [custom Python code](https://github.com/Neubias-WG5/neubiaswg5-utilities)

NetMets metrics: Geometric False Negative rate (FNR), Geometric False Positive rate (FPR) computed by [NetMets Python code](https://github.com/Neubias-WG5/neubiaswg5-utilities/blob/master/neubiaswg5/metrics/netmets_obj.py)

Metrics parameters:

GATING_DIST (UVR): Maximum distance between skeleton voxels in reference and prediction skeletons to be considered as matched (default = 5 pix)

Sigma (NetMets): tolerance in centreline position (default: 5 pix)

**Problem class: Filament Networks Tracing (LooTrc)**

Task: Estimate the medial axis of one or several connected filament network(s)

Object Encoding: 2D/3D skeleton binary masks with skeleton pixels > 0, background = 0

Reported metrics:

UNMATCHED_VOXEL_RATE (UVR), computed by [custom Python code](https://github.com/Neubias-WG5/neubiaswg5-utilities).

NetMets metrics: Geometric False Negative rate (FNR), Geometric False Positive rate (FPR) computed by [NetMets Python code](https://github.com/Neubias-WG5/neubiaswg5-utilities/blob/master/neubiaswg5/metrics/netmets_obj.py)

Metrics parameters:

GATING_DIST (UVR): Maximum distance between skeleton voxels in reference and prediction skeletons to be considered as matched (default = 5 pix)

Sigma (NetMets): tolerance in centreline position (default: 5 pix)

Skeleton sampling distance (NetMets): skeletons are sampled to be converted to SWC models. (default: 3 voxels, default Z Ratio: 1)

**Problem class: Landmark Detection (LndDet)**

Task: Estimate the position of specific feature points

Object Encoding: 2D/3D class masks, exactly 1 landmark per non null pixel, gray level encodes landmark class (1 to N, N is the number of landmarks)

Reported metrics: Number of reference / predicted landmarks (NREF, NPRED), Mean distance from predicted landmarks to closest reference landmarks with same class (MRE). All metrics computed by [custom Python code](https://github.com/Neubias-WG5/neubiaswg5-utilities)

**Problem class: Particle Tracking (PrtTrk)**

Task: Estimate the tracks followed by particles (no division)

Object Encoding: 2D/3D label masks, exactly 1 particle per non null pixel, gray level encodes particle track ID

Reported metrics: Normalized pairing score alpha (NPSA), Full normalized pairing score beta (FNPSB), Number of reference tracks (NRT), Number of candidate tracks (NCT), Jaccard Similarity Tracks (JST), Number of paired tracks (NPT), Number of missed tracks (NMT), Number of spurious tracks (NST), Number of reference detections (NRD), Number of candidate detections (NCD), Jaccard similarity detections (JSD), Number of paired detections (NPD), number of missed detections (NMD), Number of spurious detections (NSD)

All metrics computed by [Particle Tracking Challenge](http://bioimageanalysis.org/track/) Java code ([archived here](https://github.com/Neubias-WG5/neubiaswg5-utilities/blob/master/bin/TrackingPerformance.jar))

Metrics parameters: GATING_DIST (default = 5, maximum distance between particle detections in reference / prediction tracks to be considered as matching)

**Problem class: Object Tracking (ObjTrk)**

Task: Estimate object tracks and segmentation masks (with possible divisions)

Encoding: 2D/3D TIFF label masks, gray level encodes object ID + division text file (see [Cell Tracking Challenge](http://celltrackingchallenge.net/datasets/) format)

Reported metrics: Segmentation measure (SEG), Tracking measure (TRA). All computed from [Cell Tracking Challenge](http://celltrackingchallenge.net/evaluation-methodology/) metric command-line executables (archived [here](https://github.com/Neubias-WG5/neubiaswg5-utilities/blob/master/bin/SEGMeasure) and [here](https://github.com/Neubias-WG5/neubiaswg5-utilities/blob/master/bin/TRAMeasure))

### Supplementary Section 6. Additional features

This section describes additional tools, different ways to interact with a BIAFLOWS server, and content migration / importation.

**Using BIAFLOWS as an image source for a Jupyter notebook**

The procedure is self-documented online, press
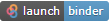
 from this GitHub repository: <https://github.com/Neubias-WG5/biaflows_jupyter_minimal>

**Importing existing datasets from a BIAFLOWS instance**

Content migration from an existing instance (e.g. BIAFLOWS online instance) to a local instance is possible. To migrate data, we developed tools that rely on BIAFLOWS RESTful programming interface to export project data (including images and object annotations) from the source instance to the destination instance. Corresponding code and documentation is available here and can be adapted for specific purposes:

<https://github.com/Neubias-WG5/Cytomine-project-migrator>.

**Manual Import of annotations**

In order to be able to compute metrics for benchmarking (optional), ground-truth annotations should also be provided for all uploaded images and encoded with format specified in Supplementary Section 5. Workflow results use the same formats as ground truth annotations, but to be visualized in BIAFLOWS they are internally converted to polygons and points. It is possible to manually convert annotations from an annotation image mask (or CSV files) to this format with the Python library: <https://github.com/Neubias-WG5/neubiaswg5-utilities/blob/master/neubiaswg5/helpers/data_upload.py>.Supported formats include 2D, 3D/2D+t and 3D+t objects. These annotations can then be uploaded to BIAFLOWS and displayed using overlays in the web image viewer as any annotation automatically created by a workflow.

**RESTful API documentation**

All interactions with BIAFLOWS are performed through a RESTful API. This API enables communication between a BIAFLOWS instance and a client. All the services provided by the API are summarized in a user-friendly way on a website automatically installed with BIAFLOWS installation procedure. From this interface, the documentation can be browsed and API queries / replies directly tested in a playground area:

BIAFLOWS RESTful API documentation can be found here:

[https://biaflows.neubias.org/restApiDoc/?doc_url=https://biaflows.neubias.org/restApiDoc/api#](https://biaflows.neubias.org/restApiDoc/?doc_url=https://biaflows.neubias.org/restApiDoc/api)

and here from a local instance of BIAFLOWS:

<http://biaflows/restApiDoc/?doc_url=http://biaflows/restApiDoc/api>

**Executing a BIAFLOWS workflow without BIAFLOWS server**

It is possible to run a workflow image independently of any BIAFLOWS server. This can for instance be useful to process a local folder of images. For this, first install [Docker](https://docs.docker.com/) on the target workstation, then:

- Get the docker image of the workflow from Dockerhub:
  docker pull {remote_image}

Or, alternatively, build workflow Docker image from source (GitHub repository)
Inside repository folder: docker build -t {local_image} .

- Prepare an empty folder {DATA_PATH} with a subfolder **/data** and subfolders:
  - **{DATA_PATH}/data/in**: add input images to this folder*****
  - **{DATA_PATH}/data/out**: workflow results are exported to this folder
  - **{DATA_PATH}/data/gt**: leave empty

***** Images should be 8/16-bit TIFF (2D) or 8/16-bit single file OME-TIFF (C,Z,T).

The string **_lbl** is forbidden in image name since it is used to identify ground truth annotation images.

- Run the workflow with the local flag:

docker run -v {DATA_PATH}/data:/data -it {image_name} {WORKFLOW_PARAMETERS} --infolder /data/in --gtfolder /data/gt --outfolder /data/out --local

This whole procedure is illustrated in the following Python Jupyter notebook:

<https://github.com/Neubias-WG5/biaflows_jupyter_local>

**Notes**:

--local (-l): do not download nor upload any content from / to BIAFLOWS. The images (input and ground truth) are read from specified folders. Metrics are optionally displayed to standard output.

For a more fine-grained control over BIAFLOWS interactions:

--no_download (-nd): images and ground truth are not downloaded from BIAFLOWS

--no_annotations_upload (-nau): annotations are not uploaded to BIAFLOWS

--no_metrics_computation (-nmc): metrics are not computed

--no_metrics_upload (-nmu): metrics are not uploaded to BIAFLOWS.

### Supplementary Section 7. Benchmarking Metrics

In this Section we describe benchmarking metrics as computed by BIAFLOWS. Code is available from our metrics library: <https://github.com/Neubias-WG5/neubiaswg5-utilities>.

Object Segmentation (ObjSeg)

**DICE (DICE)**

DICE coefficient is computed as:

**2 *** **AreaOverlap**( ***X***, ***Y*** ) / ( **Area**( ***X*** ) + **Area**( ***Y*** ) )

where ***X*** is ground truth binary mask and ***Y*** is prediction binary mask*.* Object pixels are nonnull pixels in the original masks.

DICE coefficient ranges from 0 (no overlap between segmented objects) to 1 (perfect overlap).

**AVERAGE HAUSDORFF DISTANCE (AHD)**

The Average Hausdorff Distance (AHD) is:

(Avg( **D_1_** ) **+** Avg( **D_2_** )) / 2

where, for every object pixel of the ground truth mask the minimum distance **D_1_** to the closest object pixel of the prediction mask is computed (and vice versa, leading to minimum distance **D_2_** for every object pixel of the prediction mask). Object pixels are all nonnull pixels of the masks, and averages are computed over the object pixels of the respective masks. Lower AHD implies better segmentation and the metric is equal to 0 only for perfect segmentation.

**FRACTION OVERLAP**

Every object **R** from the ground truth label mask is associated to the object **P** from the prediction label mask with maximum overlap. The score for this object is computed as:

**S** = **AreaOverlap** ( **R**, **P** ) **/ max**( **Area** (**R**), **Area** (**P**) )

The reported metric is the average score **S** over all objects **R**.

A 0.5 fraction overlap can be interpreted as "on average the area of a predicted object overlapping with the ground truth object with largest overlap is half the area of the larger of these two objects". The metric is equal to 1 only for perfect segmentation. This would for instance happen if an object is erroneously split into two objects of the same size, or if two objects of the same size are erroneously merged.

Mean Average Precision

Mean average precision as used in Data Science Bowl 2018 evaluation: https://www.kaggle.com/c/data-science-bowl-2018/overview/evaluation

Intersection over union (**IoU**) between two sets of pixels is calculated as:

**IoU(A,B) = (A ∩ B) / (A ∪ B)**.

The metric calculates IoU between all predicted objects and ground-truth objects. Then IoU for each predicted object is tested with 10 thresholds (0.5, 0.55, ..., 0.95) and if the IoU is greater than the tested threshold, the object is set as true positive (**TP**) for that threshold.

Precision at each threshold value is calculated as

**P = TP / (TP + FP + FN)**,

where **FP** = number of predicted objects - TP and **FN** = number of ground-truth objects - TP.

Average precision (**AP**) of a single image is the mean of precision with 10 different thresholds.

Mean average precision (**mAP**) is the mean of AP for all images.

Spot / object counting (ObjCnt)

**RELATIVE_ERROR_COUNT (REC)**

The relative error count is computed as the absolute difference between the number of ground truth and prediction objects, normalized to the number of ground truth objects. A perfect count leads to REC = 0.

Spot / object detection (ObjCnt)

**CONFUSION_MATRIX (TP, FN, FP)**

**F1_SCORE (F1)**

**PRECISION (PR)**

**RECALL (RE)**

**Distance RMSE (RMSE)**

The set of ground truth and prediction detections are first associated by solving a distance-constrained assignment problem with the Hungarian algorithm. This procedure pairs detections up to a maximum (gating) distance by minimizing their overall distance. Paired detections are considered True Positives (**TP**). In a classification setup, given a classification problem with ***C*** distinct classes, a confusion matrix is a square matrix of dimensions ***C*** x ***C*** where the element ***a_ij_*** (***i*** ∈ [0, ***C***[, ***j*** ∈ [0, ***C****[*) is the number of samples of class ***j*** that are predicted to be class ***i***. For binary classification (***C*** = 2), ***a_10_*** is the number of False Positives (**FP**) and **a_01_** is the number of False Negatives (**FN**).

The precision is defined as (see definitions of True Positive and False Positive metrics):

PR = TP / (TP + FP)

The precision reflects the ability of a classifier not to label as positive a sample that is negative. It ranges from 0 (all positive samples are misclassified) to 1 (no false positive). For Multiclass classification (*C* > 2), we use the weighted averaged precision: precision is computed for each class separately and then the resulting precisions are averaged weighted by support (number of positive samples for each class). This weighting strategy accounts for class imbalance.

The Recall is defined as (see definitions of True Positive and False Negative metrics):

RE = TP / (TP + FN)

Intuitively, the recall is the ability of the classifier to find all positive samples. It ranges from 0 (all positive samples misclassified) to 1 (no false negative). For Multiclass classification, we use the same approach as for Precision (**PR**): a support weighted average recall.

The accuracy (ACC) is the proportion of correctly predicted samples (the sum of the diagonal elements of the confusion matrix divided by the total number of samples). The accuracy ranges from 0 (all samples misclassified) to 1 (all samples correctly classified). A classifier that would pick a class at random typically yields accuracy around 1 / **C**. For binary classification, we have:

ACC = (TP + TN) / (TP + TN + FP + FN)

The F1-score is defined as:

F1 = 2 * (PR * RE) / (PR + RE)

where **PR** is the Precision and **RE** is the Recall (see the definition of these metrics). Recall and precision must be analyzed jointly and it makes no sense to benchmark one or the other independently. For instance, for a balanced binary classification problem, it is easy to get a perfect recall of 1 with a constant classifier that would always predict the positive class but this classifier would yield a precision of 0.5 (which is the score a random classifier would get). This is what F1-score attempts to capture. It ranges from 0 (precision and / or recall equal 0) to 1 (all samples correctly classified).

Localization accuracy (root mean square distance) of paired detections between ground truth and predictions. The default gating distance (maximum pairing distance) is set to 5 pixels. For instance, a RMSE distance of 3 means that on average the detected particles (objects) are 3 pixel away from the ground truth particles (objects).

Pixel/Voxel Classification (PixCla)

**F1_SCORE (F1)**

**ACCURACY (ACC)**

**PRECISION (PR)**

**RECALL (RE)**

See definitions from Spot / object detection (ObjCnt)

Landmark Detection (LndDet)

Number of ground truth landmarks (NREF)

Number of predicted landmarks (NPRED)

Mean distance (MRE): Mean distance from predicted to ground truth landmarks (pix)

Particle Tracking (PrtTrk)

14 metrics from [PTC challenge](https://media.nature.com/original/nature-assets/nmeth/journal/v11/n3/extref/nmeth.2808-S1.pdf) (supplementary note 3), of which 5 metrics are derived:

1. α(X, Y) = 1−d(X, Y)/d(X, Ø). Ø denotes a set of dummy tracks; hence, d(X, Ø) is the maximum possible total distance (error) from the ground truth. The measure ranges from 0 (worst) to 1 (best), indicating the overall degree of matching of ground truth and predicted tracks without taking into account spurious (non-paired estimated) tracks.

2. β(X, Y) = (d(X, Ø)−d(X, Y))/(d(X, Ø) + d(Y, Ø)). Y denotes the set of spurious tracks, and d(Y, Ø) is the corresponding penalty term. The measure ranges from 0 (worst) to α (best) and is essentially α with a penalization of non-paired estimated tracks.

3. JSC = TP/(TP + FN + FP). This is the Jaccard similarity coefficient for track points. It ranges from 0 (worst) to 1 (best) and characterizes overall particle detection performance. TP (true positives) denotes the number of matching points in the optimally paired tracks; FN (false negatives), the number of dummy points in the optimally paired tracks; and FP (false positives), the number of non-matching points including those of the spurious tracks.

4. JSCθ = TPθ/(TPθ + FNθ + FPθ). This is the Jaccard similarity coefficient for entire tracks instead of single track points. Similarly to JSC, it ranges from 0 (worst) to 1 (best). TPθ denotes the number of predicted tracks paired with ground-truth tracks; FNθ, the number of dummy tracks paired with ground-truth tracks; and FPθ, the number of spurious tracks.

5. RMSE, the r.m.s. error, indicates the overall localization accuracy of matching points in the optimally paired tracks (the TP as in JSC), being a nonnegative number with the upper bound given by the maximum distance specified.

Note: gating distance to pair particles from ground truth and prediction default to 5 pixels.

Object Tracking (ObjTrk)

The metrics are computed from Cell Tracking Challenge: <http://celltrackingchallenge.net/> code and are described in detail in: <https://doi.org/10.1038/nmeth.4473>.

Segmentation accuracy measure (SEG) evaluates the average amount of overlap, being expressed by the Jaccard similarity index, between the ground truth segmentation ground truth and the prediction. Tracking accuracy measure (TRA) is a normalized weighted distance between the prediction and the ground truth, with weights chosen to reflect the effort it takes a human curator to carry out the edits manually (see <https://doi.org/10.1371/journal.pone.0144959> for details).

Both SEG and TRA take values in the interval [0, 1], with higher values corresponding to better performance.

Filament Networks Tracing (LooTrc)

**NetMets** metrics

Geometric False Negative rate (*G_FNR_*)

Geometric False Positive rate (*G_FPR_*)

Unmatched voxel rate (UVR)

Citations: Mayerich, D., Bjornsson C.,Taylor J., Roysam B. (2012). NetMets: software for quantifying and visualizing errors in biological network segmentation. BMC Bioinformatics, 13(Suppl 8): S7.

NetMets relies on mapping nodes between ground truth and prediction filament networks. Given two networks **N_1_** and **N_2_**, it estimates the integrated length of sections of **N_1_** without correspondence in **N_2_** normalized to **N_1_** length. This is estimated by surrounding **N_2_** by a penalty weighting Gaussian envelope (penalty increase with distance) and integrating along **N_1_**. In order to estimate both missed and erroneously detected sections, a bi-directional measurement is performed by inverting the role of the networks: this leads to G_FNR_  and G_FPR_.

Overall, the more distant nodes are from the nearest node in the other network, the lower the metrics (1: perfect match).

Note: The skeleton masks are first converted to SWC models by sampling them (default sampling step is 3 voxels and default Z ratio is 1).

**Unmatched voxel rate (UVR)**

Ground truth and prediction skeleton masks are both dilated by a gating distance (default: 5 voxels). The number of object voxels from prediction mask not falling within the gating distance of an object voxel of the ground truth are counted. The same is performed for ground truth object voxels (respect to prediction object voxels). Unmatched Voxel Rate (UVR) is computed as the number of unmatched voxels (two-ways) divided by the sum of the number of object voxels in both skeleton mask. It equals to 0 for perfect matching (within the gating distance) and to 1 for the worse possible matching.

Filament Tree Tracing (TreTrc)

Geometric False Negative rate (*G_FNR_*)

Geometric False Positive rate (*G_FPR_*)

See metrics definition from Filament Networks Tracing (LooTrc). The note does not apply since workflows directly outputs the trees in SWC format.

1. Used features: Gaussian Smoothing, Laplacian of Gaussian, Gaussian Gradient Magnitude, Difference of Gaussians, Structure Tensor Eigenvalues, Hessian of Gaussian Eigenvalues. All with sigma 0.70 and 1.00.
2. SGD optimizer, mask_rcnn_presegmentation.h5 described in [5]. Backbone initialized with Coco pre-trained weights. Training, 1 epoch all layers with lr = 0.001, 1 epoch 5+ layers with lr = 0.0005, 1 epoch head layers with lr = 0.0001. Loss function as defined in [1] (mean binary cross-entropy).
3. lr = 1e^-4^, 15 epochs (500 steps), batch size: 10, loss: weighted cross-entropy, optimizer: RMSprop.
